# DeepMicroClass sorts metagenomes into prokaryotes, eukaryotes and viruses, with marine applications

**DOI:** 10.1101/2021.10.26.466018

**Authors:** Shengwei Hou, Tianqi Tang, Siliangyu Cheng, Ting Chen, Jed A. Fuhrman, Fengzhu Sun

**Author notes:** Correspondence (S. Hou) and (F. Sun). These authors contributed equally to this work (Shengwei Hou, Tianqi Tang, Siliangyu Cheng).

## Abstract

Sequence classification reduces the complexity of metagenomes and facilitates a fundamental understanding of the structure and function of microbial communities. Binary metagenomic classifiers offer an insufficient solution because environmental metagenomes are typically derived from multiple sequence sources, including prokaryotes, eukaryotes and the viruses of both. Here we introduce a deep-learning based (as opposed to alignment-based) sequence classifier, DeepMicroClass, that classifies metagenomic contigs into five sequence classes, i.e., viruses infecting prokaryotic or eukaryotic hosts, eukaryotic or prokaryotic chromosomes, and prokaryotic plasmids. At different sequence lengths, DeepMicroClass achieved area under the receiver operating characteristic curve (AUC) scores >0.98 for most sequence classes, with the exception of distinguishing plasmids from prokaryotic chromosomes (AUC scores *≈* 0.97). By benchmarking on 20 designed datasets with variable sequence class composition, we showed that DeepMicroClass obtained average accuracy scores of ∼0.99, ∼0.97, and ∼0.99 for eukaryotic, plasmid and viral contig classification, respectively, which were significantly higher than the other state-of-the-art individual predictors. Using a 1-300 µm daily time-series metagenomic dataset sampled from coastal Southern California as a case study, we showed that metagenomic read proportions recruited by eukaryotic contigs could be doubled with DeepMicroClass’s classification compared to the counterparts of other alignment-based classifiers. With its inclusive modeling and unprecedented performance, we expect DeepMicroClass will be a useful addition to the toolbox of microbial ecologists, and will promote metagenomic studies of under-appreciated sequence types.

## Introduction

Microbes are major players of global biogeochemical cycles owing to their high abundance, immense diversity, versatile metabolism, and survivability in any conceivable ecosystem on the planet (Falkowski et al., 2008; Azam & Worden, 2004). Microbial communities are a collection of diverse biological entities, including ribosome-encoding cellular organisms (REOs), capsid-encoding organisms (CEOs, i.e., viruses) that can only reproduce within cells of REOs, and orphan replicons (plasmids, transposons, etc) that parasitize REOs or CEOs for propagation (Raoult & Forterre, 2008). Viruses and plasmids are extrachromosomal genetic elements that have important implications for the diversity and function of microbial communities owing to their roles in transferring genetic materials between or within microbes. Thus, together with transposable elements, they are collectively referred to as mobile genetic elements (MGEs). Depending on where, when and how metagenomic samples were collected, the microbial diversity within a sample can range from a consortium of several dominant strains to a conglomerate of thousands of species. Soon after the discovery of the small subunit rRNA gene (SSU) as a universally conserved phylogenetic marker (Woese & Fox, 1977), the biodiversity and structure of environmental microbial communities can be easily assessed using the SSU-based amplicon surveys (Pace et al., 1986; Olsen et al., 1986). Microbial coding potentials can be further probed using cloning libraries of natural microbial assemblages (e.g., cosmid and fosmid libraries) (Olsen et al., 1986; Schmidt et al., 1991; Stein et al., 1996; Vergin et al., 1998; Rondon et al., 2000; Béjà et al., 2000; Legault et al., 2006), which have been revolutionized by shotgun metagenomes to infer functional capabilities and ecological roles of uncultured microbes (Venter et al., 2004; Handelsman, 2004). The rapid expansion of metagenomic datasets presents opportunities and challenges. Metagenomics enables the exploration of complex microbial interactions and genetic evolution of individual species (Xia et al., 2011; Schloissnig et al., 2013). On the other hand, efficient and reliable retrieval of microbial genomes and MGEs from metagenomic sequence pools requires strategic approaches.

By categorizing metagenomic contigs into distinct groups, the complexity of metagenomes can be reduced to certain taxonomic levels, from coarse domains to consensus species or strains. Metagenomic applications developed to retrieve intended contigs can be briefly framed into two categories, supervised contig classification tools (i.e., viral contig predictors) and unsupervised contig clustering tools (i.e., metagenomic binners, see Sedlar et al., 2017 for a review of binning strategies). Viruses are prevalent in aquatic, soil and host-associated systems, and are presumably the most abundant biological entities on Earth (Suttle, 2005, 2007). In marine systems, viral lysis is crucial in redirecting carbon and energy flow to the lower trophic levels (termed “Viral Shunt”), which has great implications for the global biogeochemical cycles (Fuhrman, 1999; Wilhelm & Suttle, 1999). Metagenomic contig classification has been heavily focused on the prediction of viral sequences. VirSorter (Roux et al., 2015) and VirFinder (Ren et al., 2017) are two pioneer tools to identify viral contigs from metagenomic assemblies. VirSorter predicts viral contigs based on viral signals and categorizes them into three tiers with different confidence levels. VirFinder employs k-mer frequencies and logistic regression to classify contigs to either viral or host sequences, which outperforms VirSorter at different contig lengths, especially for shorter contigs without detectable viral hallmark genes (Ren et al., 2017). The success of k-mer based methods has inspired the application of deep learning in viral sequence discovery, which led to the development of DeepVirFinder (Ren et al., 2020) and PPR-Meta (Fang et al., 2019), both of which use one-hot encoding to convert DNA sequences into presence/absence matrices of nucleotides, and use neural networks to train virus-host classifiers at different contig lengths. Besides, PPR-Meta combines both nucleotide path and codon path in the encoding step, and classifies contigs into viruses, host chromosomes and plasmids (Fang et al., 2019). VIBRANT (Kieft et al., 2020) uses neural networks to distinguish prokaryotic dsDNA, ssDNA and RNA viruses based on “v-score” metrics, which are calculated from significant protein hits to a collection of Hidden Markov Model (HMM) profiles derived from public databases. Most of the aforementioned tools target bacteriophages. Eukaryotic virus predictors are emerging in recent years, and one such tool is HostTaxonPredictor (HTP) (Gałan et al., 2019), which utilizes four machine learning methods to classify viral sequences to eukaryotic viruses or bacteriophages based on sequence features including mono-, dinucleotide absolute frequencies and di-trinucleotide relative frequencies. Plasmids are another major type of MGEs heavily studied in environmental microbiome, particularly in host-associated systems or wastewater treatment plants. Via transferring among hosts or exchanging genes with their host genomes, plasmids facilitate the acquisition of new traits by hosts (Hall, 2016). Thus, by carrying genes related to resource utilization, antibiotic, metal resistance, and defense systems, plasmids contribute to the genetic and phenotypic plasticity of hosts, and increase their fitness to the changing environments. There are multiple dedicated tools developed besides PPR-Meta, such as cBar (Zhou & Xu, 2010), PlasFlow (Krawczyk et al., 2018), PlaScope (Royer et al., 2018) and PlasClass (Pellow et al., 2020). In principle, PlaScope employs a similarity searching approach based on species-specific databases, while cBar, PlasFlow and PlasClass use differential k-mer frequencies with different machine-learning methods. Beyond viruses and plasmids, there is a paucity of applications targeting the classification of eukaryotic contigs from metagenomes, while eukaryotes are indispensable to the ecological functioning of natural microbial communities. Alignment-based applications such as Kaiju (Menzel et al., 2016) and MetaEuk (Levy Karin et al., 2020) search for close matches in reference databases, thus can be used to assign reads or contigs to taxonomic groups. While the accuracy of these applications depends on the completeness of reference databases, their performance in classifying eukaryotic contigs is arguable due to the lack of a comprehensive microbial eukaryotic database (Keeling et al., 2014). EukRep (West et al., 2018) is a reference-independent application that uses k-mer frequency and linear-SVM to classify metagenomic contigs into eukaryotic and prokaryotic sequences. It has been proven that when combined with the conventional metagenomic and metatranscriptomic analyses, such as reconstructing eukaryotic bins and gene co-abundance analysis, biological and ecological insight can be readily obtained for uncultured eukaryotes (Vorobev et al., 2020; West et al., 2018). Eukaryotic sequences could also be identified using alignment-independent applications. Tiara (Karlicki et al., 2022) is a deep-learning based method used for eukaryotic sequence identification in metagenomes, and Whokaryote (Pronk & Medema, 2022) is a random forest classifier that uses gene-structure based features to distinguish eukaryotic and prokaryotic sequences.

Despite the significant progress made in the past years, there isn’t one tool that can classify eukaryotic/prokaryotic genomes, eukaryotic/prokaryotic viruses, and plasmids in one shot. In fact, all these binary classifiers suffer from sequence types that are not modeled, such as eukaryotic contigs or plasmids can be misclassified as viruses by viral predictors, and viral contigs can be misclassified as plasmids by plasmid predictors, etc. Thus, to achieve a more reliable classification of the target sequences, one has to run several of these tools consecutively, each suffers from its sensitivity and specificity, and the error rates propagate throughout the workflow, resulting in less accurate and biased classification. Here we introduce DeepMicroClass, a versatile multi-class metagenomic contig classifier based on convolutional neural networks (CNN). The implementation of DeepMicroClass and code for experiments described in this paper can be accessed at https://github.com/chengsly/DeepMicroClass. We show that DeepMicroClass outperforms all the existing tools by precision and sensitivity across all benchmark datasets with variable sequence-type composition. Using a coastal marine metagenomic dataset as a case study, we showed that DeepMicroClass captures more eukaryotic contigs than alignment-based classifiers. DeepMicroClass is superior to the other tools by classifying all sequence types simultaneously, which will greatly reduce the time and computation resource usage compared to the conventional workflow of chaining a set of different predictors.

## Materials and methods

### Dataset preparation

We collected 5 classes of sequences: prokaryotic host, eukaryotic host, plasmid, prokaryotic viral and eukaryotic viral sequences. For prokaryotic chromosome sequences, we downloaded all the prokaryotic genomes, including all the bacteria and archaea sequences from NCBI RefSeq on Aug 22, 2022. The prokaryotic genomes were cleaned up by removing all the sequences annotated as “Plasmid” according to the assembly reports, and sequences not annotated as plasmid but have identical sequence IDs in the plasmid dataset were also removed. The resulting sample set contains 40,208 sequences. The eukaryotic host sequence database includes eukaryotic sequences from the eukaryotic taxa used by Kaiju (Menzel et al., 2016) and the PR2 database (Guillou et al., 2013). Specifically, we selected microbial eukaryotic genomes under taxa names: “Amoebozoa”, “Apusozoa”, “Cryptophyceae”, “Euglenozoa”, “Stramenopiles”, “Alveolata”, “Rhizaria”, “Haptista”, “Heterolobosea”, “Metamonada”, “Rhodophyta”, “Chlorophyta”, and “Glaucocystophyceae” using genome_updater (available at https://github.com/pirovc/genome_updater) on Aug 22, 2022. A total of 612 eukaryotic sequences were downloaded. In addition to these eukaryotic genomes, we also included 32,073,625 eukaryotic host sequences from the 678 marine eukaryotic transcriptomic re-assemblies (Johnson et al., 2019) of cultured samples generated by the MMETSP project (Keeling et al., 2014), which included 306 pelagic and endosymbiotic marine eukaryotic species representing more than 40 phyla.

Plasmid sequences and corresponding metadata were retrieved from PLSDB (Galata et al., 2019) released on Jun 23, 2021. The dataset contains 34,513 plasmid records. Viral sequences and associated metadata were retrieved from Virus-Host DB (Mihara et al., 2016) released on Jun 1, 2022, which contains 17,357 nucleic acid records, including 5,209 prokaryotic viruses and 12,148 eukaryotic viruses. In all downloaded sequences, we further cross compared sequence IDs in each class, and any sequence with an identical ID occurring in more than one class was removed so that we could reduce potential erroneous annotation from the source database.

### Benchmark Dataset Preparation

Sequences were split into two parts according to the dates submitted to NCBI, using Jan 1, 2020 as a cutoff date. Sequences submitted before Jan 1, 2020 were used for training and validation, with 80% as training and 20% as validation using stratified split, and the sequences submitted after this date were used for testing. The Mash (Ondov et al., 2016) distance was used to estimate the similarity between sequences among training, validation and test sets. Sequences in the test set with a Mash distance *<* 0.1 to any sequence in the training or validation sets were removed from the test set. VirusHost DB derived viral sequences (Mihara et al., 2016) and MMETSP derived eukaryotic sequences were not dated. These sequences were randomly split into training, validation and test sets with the proportions of 60%, 20% and 20%, respectively. Similarly, sequences were removed from the test set when the Mash distance *<* 0.1 to any sequence in the training or validation sets. The composition of a metagenomic sample is usually unknown, and the imbalance among different sequence classes might affect the performance of different classifiers. Moreover, existing methods focus on classifying one special sequence class, e.g. eukaryotic hosts, prokaryotic viruses or plasmids. Some tools could classify two or more sequence classes, for instance, PPR-Meta (Fang et al., 2019) can predict prokaryotic hosts, phages and plasmids. In order to compare with tools developed for a specific sequence class and for multiple sequence classes, we generated 20 equal-sized (1000 contigs, each 10 kbs long) benchmark datasets with a variable composition of the 5 sequence classes. Briefly, the fractions of PROK (including prokaryotic hosts, prokaryotic viruses, and plasmids) to EUK (including eukaryotic hosts and eukaryotic viruses) sequences were determined using the ratios of 9:1, 7:3, 5:5, 3:7, and 1:9. Then for each fixed PROK:EUK ratio, the PROK fraction was further split into prokaryotic hosts, prokaryotic viruses and plasmids based on the ratios of 5:1:1, 4:1:1, 3:1:1, and 2:1:1; and the EUK fraction was further split into eukaryotic hosts and eukaryotic viruses according to the ratio of 5:1, 4:1, 3:1, and 2:1. Finally, the corresponding number of sequences were drawn from the test sequence pool for each class using the ratios specified above, the actual sequence source composition of the 20 test datasets were shown in Fig. S1 and Table S1 in the Supplementary Material.

### Model Design and Training

DeepMicroClass employs a di-path convolutional neural network comprising a base-path and a codon-path to classify input sequences into one of the five classes. For the base-path, the input nucleotide sequence was firstly encoded as a one-hot matrix. Specifically, each of the A, C, G, and T nucleotides was translated into [1, 0, 0, 0], [0, 1, 0, 0], [0, 0, 1, 0], [0, 0, 0, 1], respectively. Any non-ACGT nucleotide was represented with [0, 0, 0, 0]. The reverse complimentary strand of the input sequence can be one-hot encoded simply by flipping the forward one-hot matrix along both row and column. For the codon-path, the forward or reverse base-path matrix was first converted into three 64 dimensional one-hot matrices based on three reading frames, then the three matrices were concatenated into one matrix. Thus, for each strand of a input contig, a di-path incorporating both the base and codon level information was encoded and fed into the following convolutional layers. The overview of the network structure of DeepMicroClass is shown in Fig. 1.

**Fig 1.**
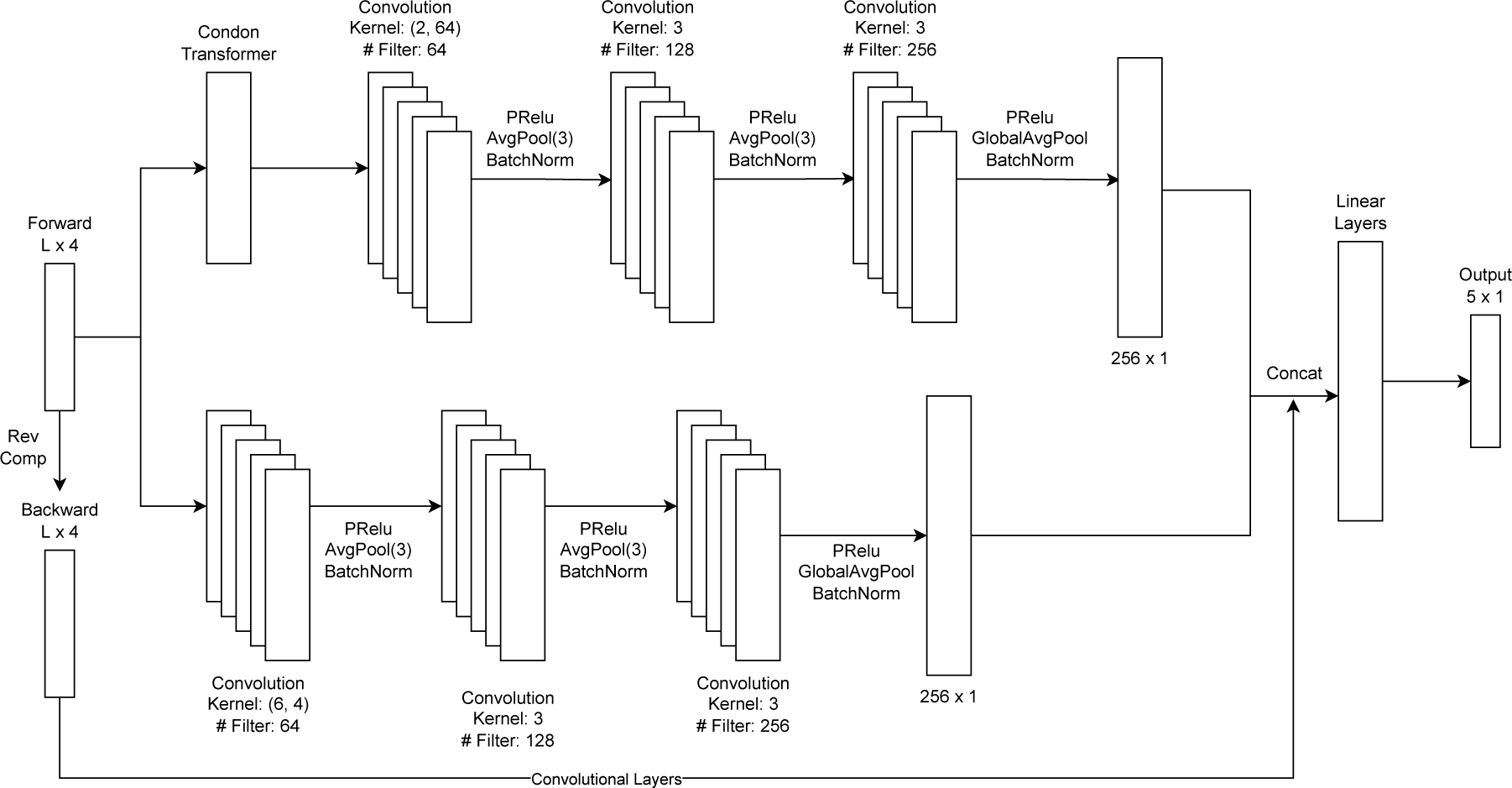
Schematic representation of the multi-class CNN structure used in this study. The network has two convolutional paths, a base-path encodes the nucleotide level information and a codon-path encodes the codon level information. The hyperparameters used for each convolutional layer are marked on the figure. For each strand, the output dimension of base- and codon-paths are 256 and 256, respectively. The di-path outputs of forward and reverse strands are concatenated into a 1024-dimensional vector, which is used as the input of following linear layers. The final linear layer outputs a 5-dimensional vector, with each dimension indicating the probability of the input contig being eukaryotic host, eukaryotic virus, plasmid, prokaryotic host and prokaryotic virus.

The di-path CNN model was trained by minimizing the cross-entropy loss between the predicted class and the actual class of input sequences. The training was run for 3000 epochs with a learning rate of 0.001 and batch size of 256. For each batch, sequences from the whole training dataset were firstly subsampled with weighted random sampling without replacement within an epoch. The weight for samples of each class *i* was defined as

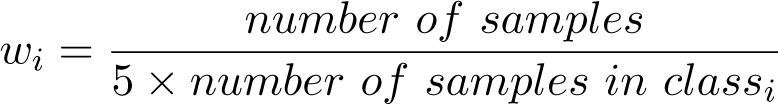

After the sequences were sampled, a contig length was chosen from 500 bps, 1 kbps, 2 kbps, 3 kbps and 5 kbps, and a contig with the given length was sampled from the original sequence to construct the batch. In the testing stage, sequences with lengths *<* 5 kbps were fed directly to the model for prediction. For sequences with lengths *>* 5 kbps, each input sequence was first split into multiple non-overlapping 5 kbps chunks, then scores given by the model for each chunk were collected, and the mean score of all chunks was used as the final output of the input sequence.

### Use-case data preparation and analysis

The daily time-series metagenomic samples were taken off the coast of Southern California using an Environmental Sample Processor (ESP), and the 1 µm A/E filters (Pall Gelman) collected during the day were used for DNA extraction as described previously (Needham et al., 2018). Metagenomic libraries were prepared using the Ovation^®^ Ultralow V2 DNA-Seq library preparation kit (NuGEN,

Tecan Genomics) under the manufacturer’s instruction using 10 ng of starting DNA and amplified for 13 PCR cycles. Metagenomic libraries were sequenced on an Illumina NovaSeq 6000 platform (2 *×* 150 bp chemistries) at Berry Genomics Co. (Beijing, China). After demultiplexing, the raw reads were first checked with FastQC v0.11.2, then adapter and low quality regions were trimmed using fastp v0.21.0 (Chen et al., 2018) with the following parameters: -q 20 -u 20 -l 30 –cut_tail -W 4 -M 20 -c. PhiX174 and sequencing artifacts were removed using bbduk.sh and human genome sequences were removed using bbmap.sh with default parameters, both scripts can be found in the BBTools package v37.24 (https://jgi.doe.gov/data-and-tools/bbtools). Metagenomic samples were assembled independently using metaSPAdes v3.13.0 (Nurk et al., 2017) with a custom kmer set (-k 21,33,55,77,99,127). The assembled contigs were further coassembled as previously described (Long et al., 2021). Briefly, all the contigs were pooled and sorted into short (*<*2kb) or long (*≥*2kb) contig sets, the short contig set was first coassembled using Newbler v2.9 (Margulies et al., 2005), the resulting *≥*2kb contigs were further coassembled with the long contig set (Treangen et al., 2011). A minimum overlap thresholds of 80 nt and 200 nt were set for Newbler and minimus2, respectively. For both coassembly steps, a minimum identity cutoff of 0.98 was applied. After co-assembly, contigs were further dereplicated at 0.98 identity using cd-hit v4.6.8 (Li & Godzik, 2006), the resulting contigs were used as reference contigs for sequence classification and read recruitment analysis. Reference contigs were classified using Kaiju v1.7.3 (Menzel et al., 2016) and MetaEuk v1 (Levy Karin et al., 2020), as well as DeepMicroClass v0.1.0 (in hybrid mode), read counts assigned to each sequence class were summarized using custom Python scripts. Reads were mapped to reference contigs using bwa mem v0.7.17 with default parameters, and the number of reads aligned *>*30 nt to reference contigs were counted using bamcov v0.1 (available at https://github.com/fbreitwieser/bamcov) with default parameters.

## Results

### A CNN-based multi-class classifier

Identifying contigs of microbial eukaryotes and the viruses infecting them from metagenomic assemblies is crucial for gaining a better understanding of their ecological roles. However, current state-of-the-art tools often do not fully appreciate most of the eukaryotic viruses and their hosts. Here two commonly used viral contig predictors, VirFinder (Ren et al., 2017) and PPR-Meta (Fang et al., 2019), were evaluated based on their predicted viral scores. As expected, both predictors gave high scores to prokaryotic viral sequences and low scores to prokaryotic host sequences. However, the scores for eukaryotic host and eukaryotic viral sequences were more evenly distributed (**Fig. S2**), revealing an insufficient accuracy in classifying these sequence classes. Out of 500 randomly subsampled genomic sequences for each sequence type of prokaryotes, prokaryotic viruses, microbial eukaryotes, and eukaryotic viruses downloaded from NCBI, 454 prokaryotic viruses and 85 prokaryotic hosts had VirFinder-scores (VFscores) above 0.5, while 238 eukaryotic viruses and 157 eukaryotic hosts had VF-scores above this value (**Fig. S2a**). A similar trend can be observed for PPR-Meta (**Fig. S2b**), confirming these tools are not adequately equipped to handle eukaryotic viral and host sequences. This emphasizes the need for novel predictors that consider more sequence types during the model training process.

Here the performance of DeepMicroClass on sequences with different lengths (500 bps, 1 kbps, 2 kbps, 3 kbps, 5 kbps, 10 kbps, 50 kbps, and 100 kbps) was evaluated on test data. The model performance for each sequence type was visualized via the Receiver Operating Characteristics (ROC) curve using a one-versus-rest strategy (**Fig. 2**). Overall, we showed that as the sequence length increased, the model’s performance improved across most sequence types, as indicated by the Area Under the Receiver Operating Characteristic (AUC) measurements (**Fig. 2**). DeepMicroClass performed well on all sequence types when the input sequence length was *≥* 1 kbps, with the minimum AUC score being 0.963 on classifying prokaryotic sequences. At the sequence length of 500 bps, DeepMicroClass achieved fairly high AUC scores for eukaryotic (0.944) or prokaryotic (0.96) viruses, whilst the scores for both viral sequence types were always *≥* 0.99 at longer sequence lengths (*≥* 2 kbps) (**Fig. 2**). For non-viral sequences, the AUC scores were highest for eukaryotic sequences, followed by plasmid and prokaryotic genome sequences. However, a slight drop in the True Positive Rate (TPR) could be observed for eukaryotic sequences when the False Positive Rate (FPR) was near 0 (**Fig. 2**). With further investigation, the rough curve could be caused by the sharp drop in the number of available eukaryotic sequences in the training dataset, which dropped from 16,002 to 255 when the contig length changed from 10 kbps to 50 kbps.

**Fig 2.**
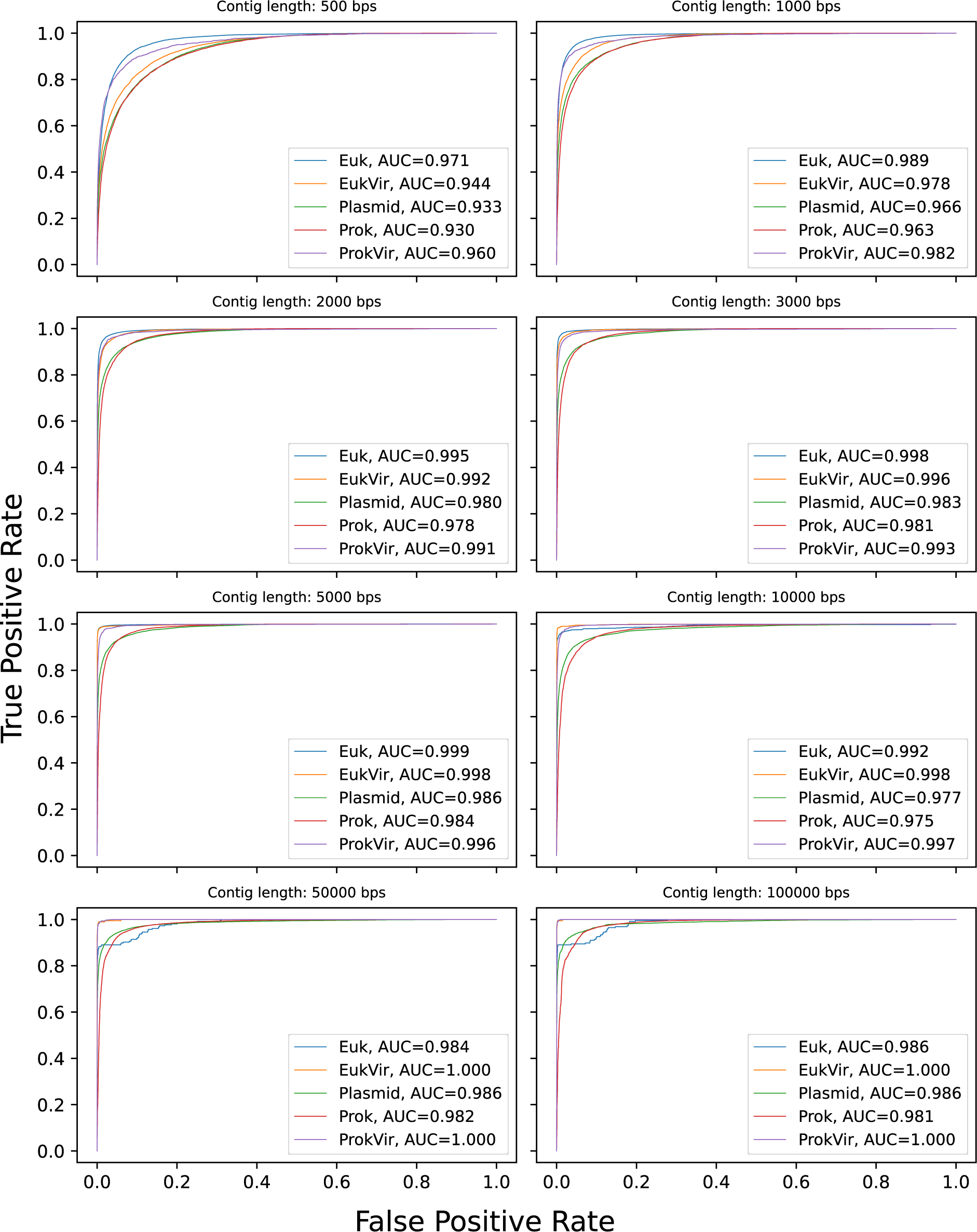
The ROC curves and AUC scores of different length models assessed on test datasets. Each different panel shows the ROC curves for 5 sequence classes at different contig lengths (500 bps, 1 kbps, 2 kbps, 3 kbps, 5 kbps, 10 kbps, 50 kbps and 100 kbps). Euk, eukaryotic sequences; EukVir, eukaryotic viral sequences; Plasmid, plasmid sequences; Prok, prokaryotic genome sequences; ProkVir, prokaryotic viral sequences.

### DeepMicroClass outperforms Tiara and Whokaryote in eukaryotic host sequence prediction

In the following three sections, we investigate the performance of DeepMicroClass for particular classes of sequences. We used accuracy and F1 score as the metrics to assess the model performance. And the sequence type composition of different benchmark datasets was described in the section Benchmark Dataset Preparation.

First, we compared the performance of DeepMicroClass with Tiara (Karlicki et al., 2022) and Whokaryote (Pronk & Medema, 2022) on the classification of microbial eukaryotes. Tiara and Whokaryote are commonly used to identify eukaryotic contigs from metagenomic assemblies without prior knowledge of microbial phylogenetic affiliation. With the compiled benchmark datasets, we showed that DeepMicroClass persistently outcompeted both tools in all scenarios in terms of accuracy and F1 score (**Fig. 3, S3**), and DeepMicroClass was robust to the different compositions of benchmark datasets (**Fig. 3**).

**Fig 3.**
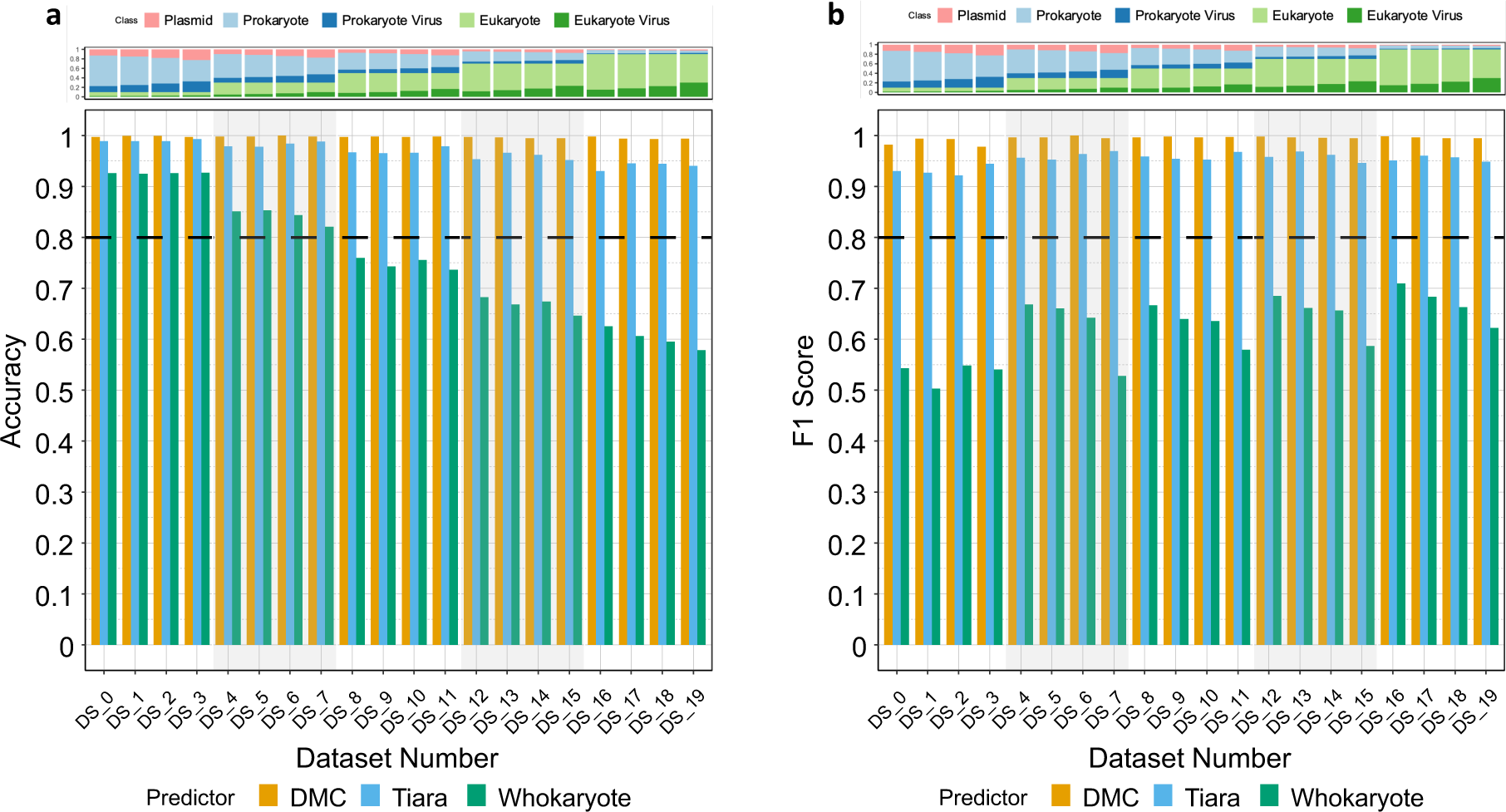
Distribution patterns of accuracy (a) and F1 score (b) across 20 benchmark datasets for DeepMicroClass, Tiara and Whokaryote. The top panel shows the sequence type composition of 20 benchmark datasets, and the detailed composition ratios can be found in Table S1. The dashed black lines indicate where accuracy or F1 score equals 0.8.

The average accuracy and F1 score across all benchmark datasets for DeepMicroClass were both 0.99, which were significantly higher than these metrics of Tiara and Whokaryote (pairwise Wilcoxon test *p*-values *≤* 9.5e-05 for both accuracy and F1 score). The accuracy of Whokaryote dropped from *∼*0.95 to *∼*0.75 as the proportion of eukaryotic sequences increased, and the F1 scores were substantially lower than 0.8 in all test datasets. In contrast, Tiara maintained high accuracy and F1 score across different eukaryotic proportions, though a slight decrease in accuracy could be observed when the eukaryotic proportion was high. DeepMicroClass achieved accuracy and F1 score above 0.98 for all tested scenarios and was robust to variable sequence composition.

A further look into those misclassified sequences revealed that both Tiara and Whokaryote suffered from lower sensitivity in distinguishing eukaryotic sequences from other types of sequences. Especially for Whokaryote, a substantial amount of eukaryotic viruses were mistakenly classified as eukaryotes (**Fig. S4**).

### DeepMicroClass outcompetes PlasFlow, PPR-Meta and geNomad in plasmid sequence classification

Plasmids are the major agents of horizontal gene transfer (HGT) among prokaryotic microbial communities. Here we compared the performance of DeepMicroClass with PlasFlow (Krawczyk et al., 2018), PPR-Meta (Fang et al., 2019) and geNomad (Camargo et al., 2023) in classifying plasmid sequences using the same benchmark datasets described above. DeepMicroClass showed significantly improved results than PlasFlow, PPR-Meta and geNomad in all tested cases in plasmid classification (pairwise Wilcoxon test adj.*p*-value *≤* 1.1e-07; **Fig. 4 & S5**). Although PlasFlow, PPR-Meta and geNomad were able to achieve a maximum F1 score of 0.68, 0.74 and 0.86, respectively, their performance was severely impaired with increasing proportions of eukaryotic sequences in the benchmark datasets (**Fig. 4**). In contrast, the F1 score of DeepMicroClass was constantly higher than 0.8, though a slight decrease could also be observed with increasing eukaryotic proportions.

**Fig 4.**
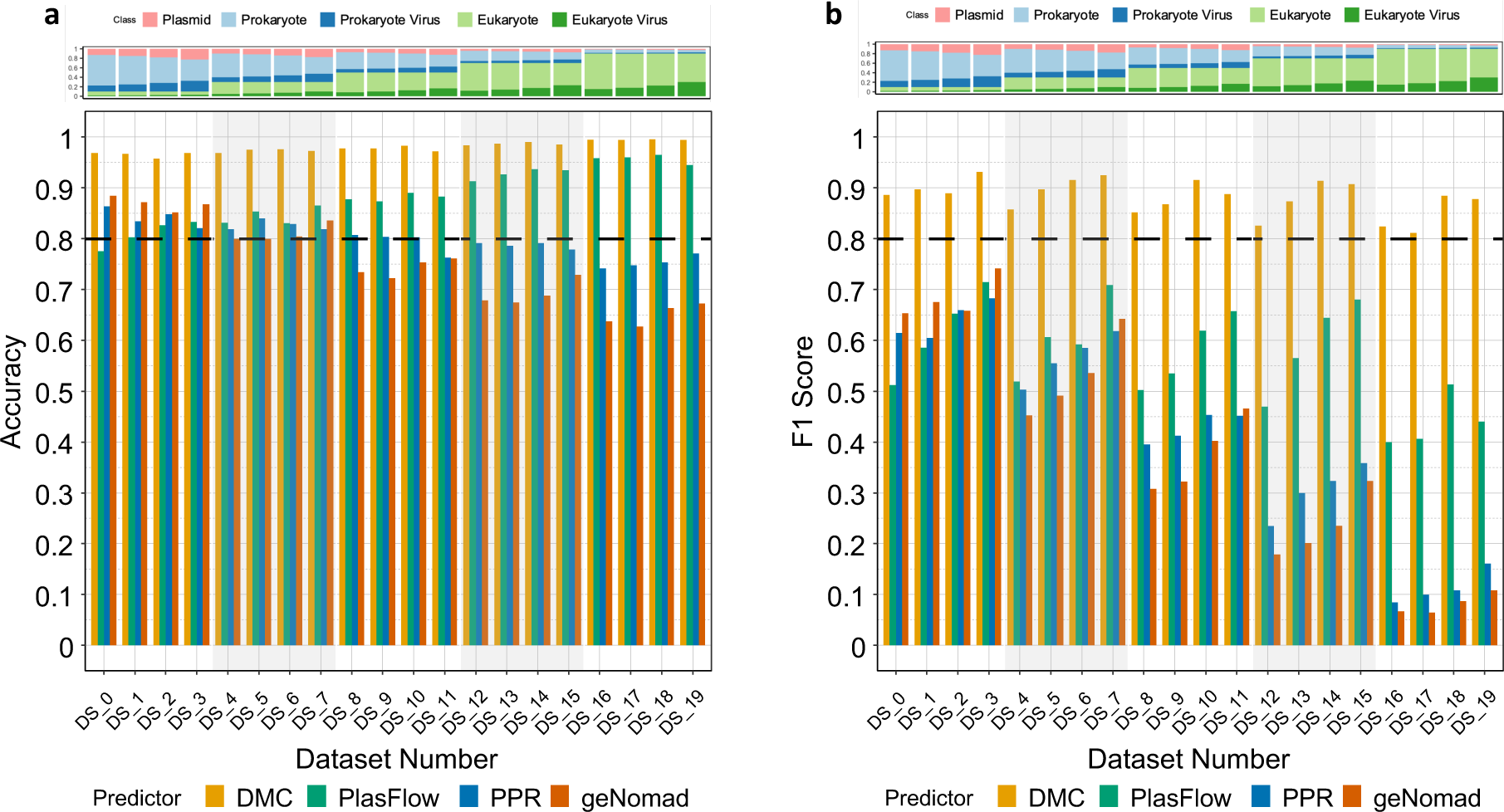
Distribution patterns of accuracy (a) and F1 score (b) across 20 benchmark datasets for DeepMicroClass, PlasFlow, PPR-Meta and geNomad on plasmid classification. The dashed black lines indicate where accuracy or F1 score equals 0.8. The same benchmarking datasets were used as in Fig. 3.

We further examined the misclassified sequences and found PlasFlow had high sensitivity but low specificity, and the dominance of misclassified sequence types was in line with the composition of benchmark datasets (**Fig. S6**). PPR-Meta might benefit from its modeling of prokaryotic chromosomes and phages, while it still had a low specificity mainly due to the misclassification of prokaryotic and eukaryotic chromosomal sequences into plasmids (**Fig. S6**). On the other hand, geNomad mainly suffered from misclassifying prokaryotic chromosomes into plasmids, though the misclassified eukaryotic sequences also accounted for a significant share (**Fig. S6**). It’s noteworthy that DeepMicroClass might further benefit from its modeling of eukaryotic genomic and viral sequences since they were rarely misclassified as plasmids, though the misclassification rates between plasmids and prokaryotic chromosomal sequences were still the highest among all misclassifications (**Fig. S12**). Probable reasons for such observation are the high affinity and frequent genetic exchange between plasmids and prokaryotic chromosomes, further improvements on the neural network structures or using additional features extracted from gene- or operon-centric approaches might yield a better classifier.

### DeepMicroClass achieves improved results in viral sequence prediction

Viruses are ubiquitously found in every natural system where cellular organisms colonize. Viral contigs have been commonly identified from metagenomes or viromes using essentially gene-centric (e.g. VirSorter (Roux et al., 2015), VirSorter2 (Guo et al., 2021), VIBRANT (Kieft et al., 2020)), or oligonucleotide-centric (e.g. VirFinder (Ren et al., 2017), DeepVirFinder (Ren et al., 2020), PPR-Meta (Fang et al., 2019)) approaches, or a combination of both approaches (e.g. geNomad (Camargo et al., 2023)). Here we compared the performance of DeepMicroClass with VirSorter2, geNomad, VIBRANT, DeepVirFinder and PPR-Meta on viral contig prediction using the aforementioned benchmark datasets. Among these methods, DeepVirFinder, VIBRANT, PPR-Meta and geNomad were trained for the prediction of prokaryotic viruses, while VirSorter2 was trained for the prediction of both eukaryotic and prokaryotic viruses. We compared the performance of DeepMicroClass with VirSorter2 on the prediction of both prokaryotic and eukaryotic viruses, and the performance of DeepMicroClass with other predictors on the prediction of prokaryotic viruses. In either case, DeepMicroClass achieved better performance in terms of both accuracy and F1 score than all the other tested tools (**Fig. 5**, **S7 & S8**). VIBRANT and VirSorter2 showed slightly lower accuracy than DeepMicroClass, followed by PPR-Meta and DeepVirFinder. More distinct differences were observed in the F1 score metric of these tools across dataset composition, DeepMicroClass achieved an average F1 score of ∼0.96, followed by VirSorter2

**Fig 5.**
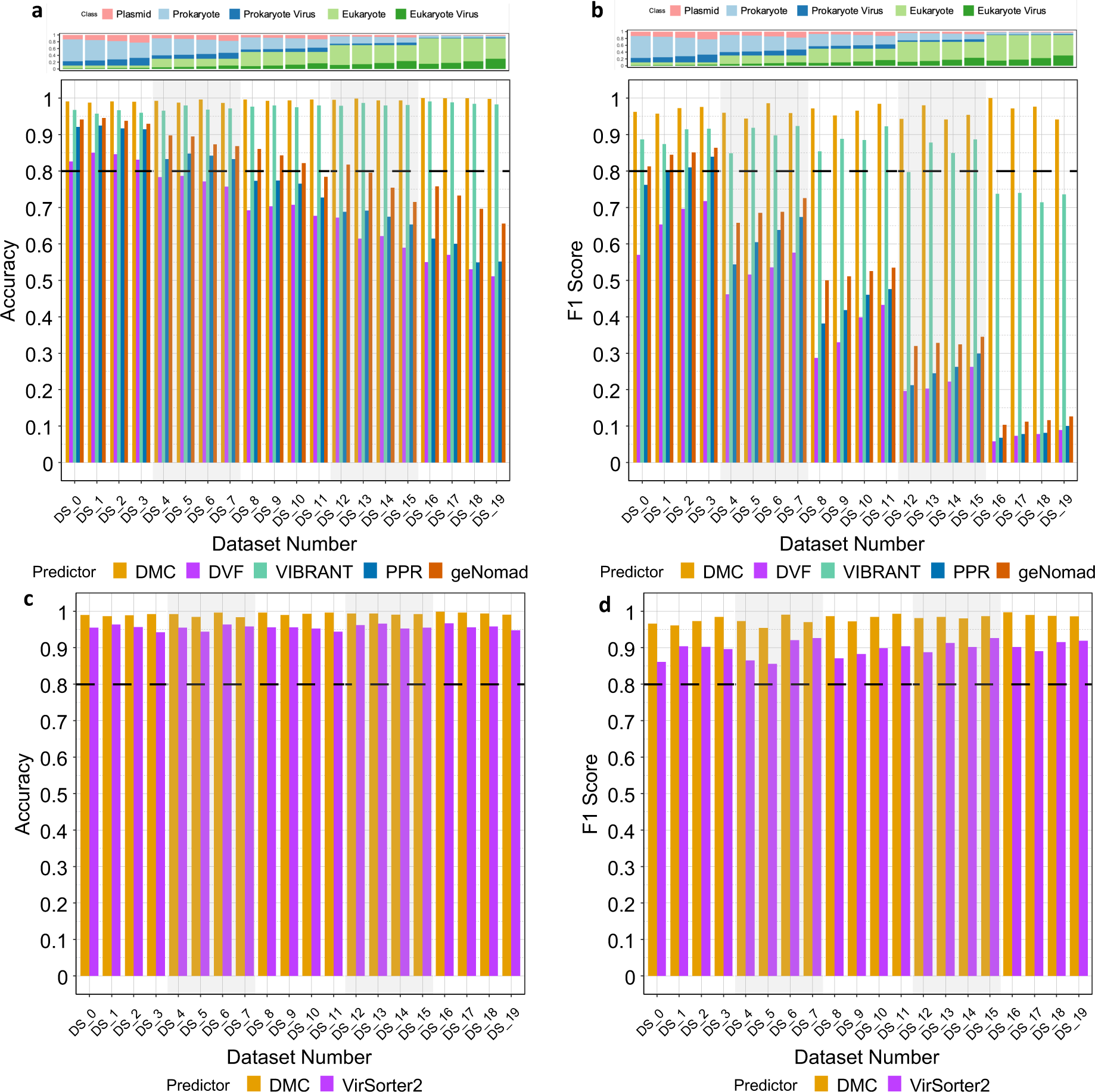
Distribution patterns of accuracy and F1 score across 20 benchmark datasets for viral classification. The accuracy (a) and F1 score (b) metrics for DeepMicroClass, PlasFlow, PPR-Meta and geNomad were evaluated on prokaryotic viral contig classification, and the accuracy (c) and F1 score (d) metrics for DeepMicroClass and VirSorter2 were evaluated on both prokaryotic and eukaryotic viral contig classification. The dashed black lines indicate where accuracy or F1 score equals 0.8. The same benchmark datasets were used as in Fig. 3. DMC, DeepMicroClass; DVF, DeepVirFinder; PPR, PPR-Meta.

DMC, DeepMicroClass; PPR, PPR-Meta and VIBRANT (∼0.90 and ∼0.85, respectively). The F1 score of VIBRANT dropped from 0.94 to <0.80 as increasing proportions of eukaryotic chromosomal and viral sequences in the benchmark datasets. PPR-Meta and DeepVirFinder showed a decreasing tendency in both accuracy and F1 score with the increasing of eukaryotic chromosomal and viral sequences (**Fig. 5a & 5b, S7**). When considering both prokaryotic and eukaryotic viral sequences as the positive viral set, DeepMicroClass and VirSorter2 were both able to achieve accuracy >0.90 and F1 score >0.80 without being significantly affected by the variations of sequence type composition, and DeepMicroClass constantly outperformed VirSorter2 in both metrics across the benchmark datasets (**Fig. 5c & 5d**, **S8**).

The number of misclassified sequences by PPR-Meta, DeepVirFinder, VIBRANT, geNomad and VirSorter2 is shown in **Fig. S9**. The distribution of misclassified sequences by PPR-Meta, DeepVirFinder and geNomad showed a similar pattern, that eukaryotic chromosomal and viral sequences were prone to be misidentified as prokaryotic viruses. This indicates tools or models trained without knowledge of eukaryotic sequences are likely to behave similarly when eukaryotes are not rare in the metagenomic community. Although VIBRANT and VirSorter2 had fewer misclassified sequences compared to PR-Meta, DeepVirFinder and geNomad, both suffered from misclassifying prokaryotic chromosomal or plasmid sequences into prokaryotic viruses **Fig. S9**. Since both VIBRANT and VirSorter2 use a gene-centric approach, it’s possible that some of the viral signature genes or fragments could also be widely detected in prokaryotic genomes or plasmids as a result of frequent gene transfer among them. This contrasts with the oligonucleotide-centric tools since cross-kingdom viral infection or plasmid conjugation and gene transfer are less common.

Since DeepMicroClass, PPR-Meta and geNomad are multiclass classifiers, here we also compared their performance based on accuracy and F1 score metrics on multiclass sequence classification using the same benchmark datasets (**Fig. S10 & S11**). Here we only considered prokaryotic chromosomal, prokaryotic viral and plasmid sequences for comparison with PPR-Meta and geNomad as they were not trained for eukaryotic sequence classification. On the other hand, all five sequence types were considered for the evaluation of DeepMicroClass. In this case, DeepMicroClass still outperformed PPR-Meta and geNomad in all tested scenarios as evaluated by both the accuracy or and F1 score metrics (pairwise Wilcoxon test *p*-values *≤* 1.9e-06; **Fig. S10 & S11**). Both accuracy and F1 scores of DeepMicroClass were rarely below 0.95 across the sequence composition of the 20 benchmark datasets, while they were rarely above 0.9 for geNomad, or rarely above 0.8 for PPR-Meta (**Fig. S10**). Although the performance of DeepMicroClass was also deteriorated by the misclassification between prokaryotic chromosomal and plasmid sequences (**Fig. S12**), the amounts of misclassified sequences were significantly lower than VIBRANT, VirSorter2 or geNomad (**Fig. S9**).

### DeepMicroClass predicted more eukaryotic and viral contigs than alignment-based predictors

Alignment-based classifiers can suffer from incomplete genomic databases, particularly for complex natural environments such as marine or soil systems. To test the performance of DeepMicroClass in real metagenomic context, here we examined its performance with the other two sequence classifiers, Kaiju (Menzel et al., 2016) and MetaEuk (Levy Karin et al., 2020), using a 1-300 µm size fraction marine metagenomic dataset sampled off the coast of Southern California (Needham et al., 2018). Using the co-assembled contigs as the reference, we show DeepMicroClass classified less prokaryotic but more eukaryotic, eukaryotic viral and prokaryotic viral contigs than Kaiju and MetaEuk (**Fig. 6a**). Among all the prokaryotic contigs classified by both Kaiju and MetaEuk, 73.6% of them were predicted to be prokaryotic by DeepMicroClass, and 11.88%, 10.39%, and 4.14% of them were predicted to be eukaryotic, prokaryotic viral and eukaryotic viral sequences, respectively (**Fig. 6b**). Contigs that couldn’t be taxonomically determined by Kaiju (16.41%) or MetaEuk (10.01%) are mainly dominated by eukaryotic sequences (57.13% / 38.3%) as predicted by DeepMicroClass (**Fig. 6c & 6d**). Although MetaEuk classified more eukaryotic contigs than Kaiju (21.88% vs 15.26%, **Fig. 6a**), the latter classified more prokaryotic viral contigs (4.38% vs 1.51%, **Fig. 6a**). This is consistent with the higher percentage of prokaryotic viral sequences in the unclassified contigs of MetaEuk than Kaiju (28.86% vs 14.87%, **Fig. 6c & 6d**). By mapping reads to reference contigs, we calculated the read percentages recruited by different sequence types. The average eukaryotic read percentage recruited by DeepMicroClass (6.15%) is considerably higher than by MetaEuk (4.78%) or Kaiju (3.50%), at the expense of lower prokaryotic read percentages (13.12%, 20.60% and 20.51%, respectively, **Fig. 6f-h**). Similarly, the average read percentages of prokaryotic viral and eukaryotic viral sequences recruited by DeepMicroClass (6.07%/1.24%) are also higher than MetaEuk (0.49%/0.19%) and Kaiju (1.67%/0.37%) (**Fig. 6f-h**). Notably, though DeepMicroClass assigned less prokaryotic and more eukaryotic reads than other classifiers, the relative abundance profiles across the whole time series are highly correlated (**Fig. S13a & S13b**), and to a less extent for the prokaryotic viral read percentage profiles (**Fig. S13c**). This is not the case for eukaryotic viral read abundance profiles, where Kaiju and MetaEuk are highly correlated, but not to DeepMicroClass (**Fig. S13d**). To sum up, DeepMicroClass is more correlated with MetaEuk in eukaryotic read profiles, and more correlated with Kaiju in prokaryotic and prokaryotic viral read profiles.

**Fig 6.**
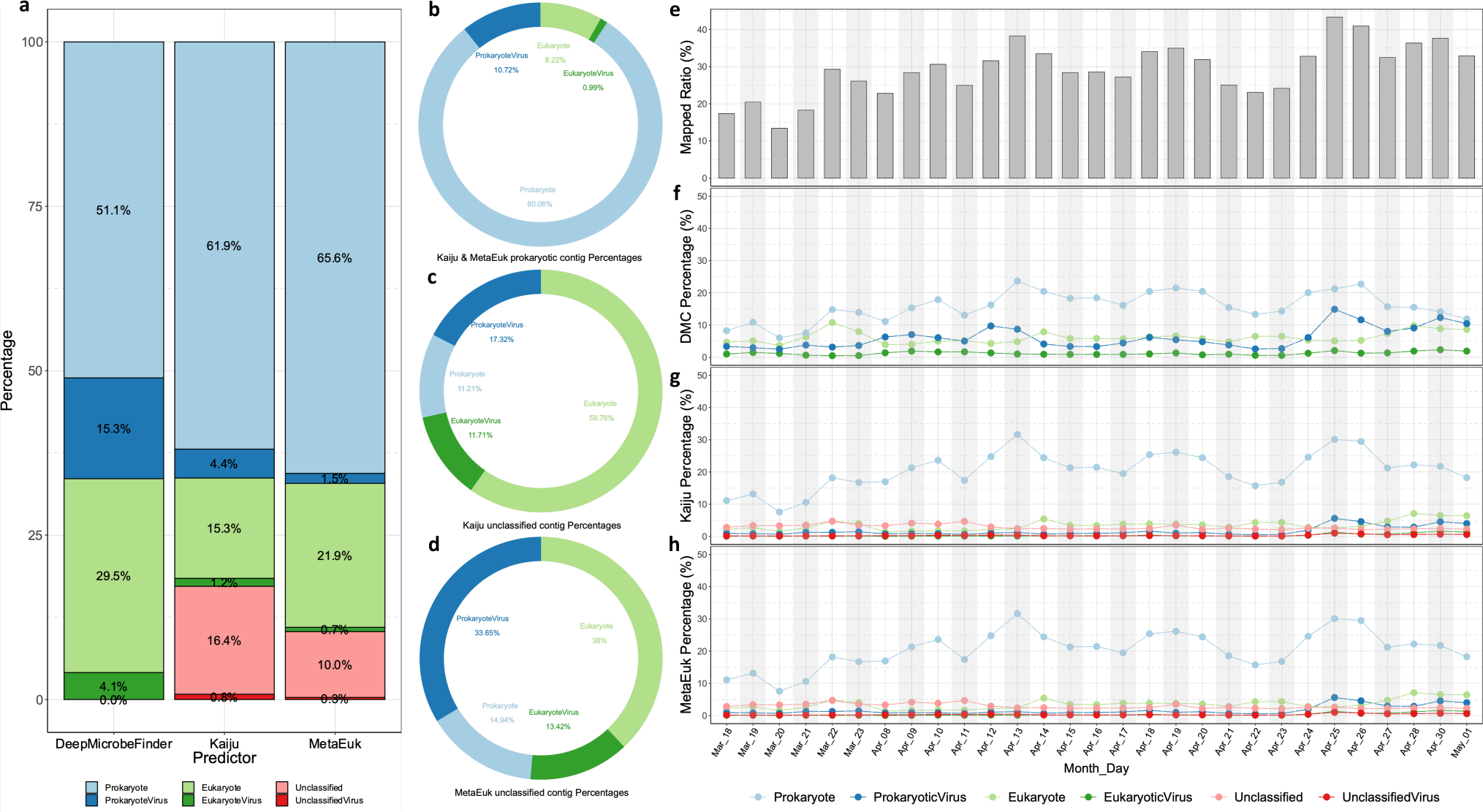
Sequence classification and read abundance of a 1-300 µm size fraction marine metagenomic dataset sampled off the coast of Southern California. Metagenomic contigs were classified using DeepMicroClass, Kaiju and MetaEuk at a length cutoff of 2 kb, and percentages of different sequence types were calculated (a). Contigs predicted as Prokaryotes by both Kaiju and MetaEuk (b), and contigs that were not classified by Kaiju (c) or MetaEuk (d) were further broken down into DeepMicroClass’s classification. Clean reads were aligned to metagenomic contigs and percentages of mappable reads were calculated (e). Mapped read percentages were further summarized according to sequence types of reference contigs as predicted by DeepMicroClass (f), Kaiju (g) and MetaEuk (h). Prokaryotes included both prokaryotic hosts and plasmids. UnclassifiedViruses were sequences predicted to be viruses but their taxonomy couldn’t be further resolved by Kaiju or MetaEuk.

## Discussion

### Microbial eukaryotes and viruses infecting them are understudied

Microbial eukaryotes are prevalent in diverse ecosystems such as host-associated habitats (Parfrey et al., 2011), deep-sea benthos (Bik et al., 2012), and geothermal springs (Oliverio et al., 2018), etc. Due to challenges in the cultivation and whole genome-sequencing of microbial eukaryotes, biodiversity surveys of microbial eukaryotes were commonly performed using marker genes, such as the 18S rDNA hypervariable V4 or V9 regions (Pawlowski et al., 2012; Amaral-Zettler et al., 2009). The ampliconbased analysis provides valuable information on the taxonomy of microbial eukaryotes, while in order to probe their metabolic potentials or ecological functions, genomic and transcriptomic information are essential. Despite several achievements in collecting microbial eukaryotic genes (Carradec et al., 2018; Vorobev et al., 2020), transcripts (Keeling et al., 2014) or single-cell amplified genomes (SAGs) (Sieracki et al., 2019) towards a comprehensive microbial eukaryotic database, our knowledge are still limited by the availability of diverse microbial eukaryotic genomes (Burki et al., 2020). With the rapid accumulation of metagenomic datasets and the availability of binning software, it’s appealing to recover eukaryotic genomes from natural microbial communities. EukRep was developed in such a context to identify eukaryotic contigs for metagenomic binning (West et al., 2018). This approach has enabled the genome-resolved analysis of fungi, protists, and rotifers from human microbiome studies (West et al., 2018; Olm et al., 2019). Similar approaches have been applied to marine microbiome studies (Duncan et al., 2020; Delmont et al., 2020), which recovered hundreds of eukaryotic metagenome-assembled genomes (MAGs) and provided insight into the functional diversity and evolutionary histories of microbial eukaryotes beyond the taxonomic information.

Beyond microbial eukaryotes, current viromic studies are biased towards viruses infecting prokaryotes. This could be introduced by the skewed distribution of viral genomes in the RefSeq database, which is dominated by phages and pathogenic viruses. By Sept 1, 2023, among 18,729 viral reference sequences, there were only 104 records belonging to algae-infecting Phycodnaviridae and 30 belonging to protists-infecting Mimiviridae. Both of the two viral families are subgroups of the Nucleocytoplasmic Large DNA Viruses (NCLDV) (Iyer et al., 2001). Since most of the commonly used viral predictors are trained on the RefSeq viral database, it’s expected that these tools suffered from identifying eukaryotic viruses from the test datasets (**Fig. 5, S7, & S8**). Given the high diversity of protists (Foissner, 1999; Slapeta et al., 2005), high throughput metagenomes and single-cell genomes are expected to offer a culture-independent solution to rapidly expand the coverage of viral database. For instance, two recent studies reconstructed 2,074 and 501 NCLDV MAGs from global environmental metagenomes (Schulz et al., 2020; Moniruzzaman et al., 2020), dramatically increased the phylogenetic and functional diversity of NCLDVs. Single-cell metagenomics was also employed to identify viruses infecting marine microbial eukaryotes (Needham et al., 2019a,b), these studies provided insightful findings of the viral encoded proteins and metabolic pathways.

These studies demonstrated that metagenomics and single-cell genomics can be promising in studying microbial eukaryotes and viruses infecting them. While most commonly used tools are not optimized in classifying eukaryotes (**Fig. 3 & S3**) or eukaryotic viruses (**Fig. 5 & S7**). Given the high performance of DeepMicroClass and the evidence of abundant eukaryotic contigs in marine ecosystems (**Fig. 6**), we expect it will be a valuable addition to the toolbox of marine ecologists.

### The challenge of classifying prokaryotic host and plasmid sequences

DeepMicroClass has a relatively lower accuracy in classifying plasmids when compared to the classification of eukaryotic or viral contigs (**Fig. 3, 4, 5**). The majority of the sequences that were misclassified as plasmids were from prokaryotic host genomes (**Fig. S12**), confirming classifying prokaryotic chromosomal and plasmid sequences is a caveat of DeepMicroClass (**Fig. 2**). In comparison, the other tested plasmid classifiers suffered from both prokaryotic and eukaryotic sequences as we have benchmarked (**Fig. 4 & S6**). It’s noteworthy that this marginal advantage can be crucial in natural environments, such as marine environments as we mentioned here (**Fig. 6**), where eukaryotic sequences can have a substantial impact on the classification of plasmid sequences. This also indicates that it is achievable to separate plasmid sequences from eukaryotic sequences solely based on patterns of oligonucleotides, and current plasmid predictors can benefit from using a more comprehensive training dataset including eukaryotic sequences.

It is understandable given the higher genome complexity of eukaryotes than prokaryotes (Lynch & Conery, 2003), such as the coding density, prevalence of introns and repetitive sequences, etc. In contrast, it’s challenging to classify plasmids and prokaryotic chromosomal sequences for all the tested plasmid predictors (**Fig. 4**). The reasons can be manifold, but plasmid transmission among microbial hosts and plasmid-chromosome gene shuffling can be two fundamental ones. The host range of plasmids is variable, it can be within closely related species for narrow host range plasmids or across distant phylogenetic groups for broad host range plasmids (Jain & Srivastava, 2013). Broad host range plasmids can be important drivers of the gene flux among host microbes in natural environments (Heuer & Smalla, 2007; Wolska, 2003; Davison, 1999). For instance, in natural soil microbial communities, the IncPand IncPromA-type broad host range plasmids could transfer from proteobacteria to diverse bacteria belonging to 11 bacterial phyla (Klümper et al., 2015). When plasmid carriage could increase the hosts’ fitness, such as improving host survival with antibiotic resistance, it can be rapidly adopted and persistently maintained in natural microbial communities (Li et al., 2020; Bellanger et al., 2014). On the other hand, when the maintenance of plasmids imposed a high fitness cost on the hosts, plasmids or plasmid-borne genes could be lost in the process of purifying selection (Hall et al., 2016). Interestingly, studies also suggested that sometimes this fitness cost could be ameliorated by compensatory evolution (Millan et al., 2014; Harrison et al., 2015; Loftie-Eaton et al., 2017), which was hypothesized to be the major factor of plasmid survival and persistence (Hall et al., 2017). Plasmid carriage also increases the chance of plasmid-chromosome genetic exchange mediated by SOS-induced mutagenesis (Rodríguez-Beltrán et al., 2021) or mobile genetic elements such as transposons and integrons, etc (Frost et al., 2005; Rodríguez-Beltrán et al., 2021). For instance, genes carried by transposons or in the variable regions were also frequently found on plasmids (Eberhard, 1990; Zheng et al., 2015). Thus, the permissive transfer of plasmids across diverse hosts and the plasmid-chromosome gene flow pose a challenge for current plasmid classifiers. The oligonucleotide-based approaches might be complemented by gene-centric approaches using plasmid signature genes or enriched gene functions, such as genes involved in mobilization or conjugation. In addition, a comprehensive plasmid database is also crucial for model training, and plasmid-enriched metagenomics (plasmidome) can be a promising way to screen plasmids from environmental samples (Shi et al., 2018).

## Conclusions

DeepMicroClass as a versatile multi-class classifier enables the accurate classification of five different metagenomic sequence types in one shot, meanwhile, it avoids the time-consuming and error-prone preprocessing steps that could potentially propagate errors to the final classification. The inclusive modeling of all common sequence types in metagenomes also makes DeepMicroClass attain better performance than the other state-of-the-art individual predictors due to reduced cross misclassifications. We also detected high relative abundances of marine eukaryotes in a daily time-series dataset, which were underestimated by alignment-based classifiers due to the limitation of public reference databases. Our case study indicates that both host and viral sequences are essential components in the cellular metagenomes, and robust ecological patterns can be obtained with DeepMicroClass even for coarse sequence types. We argue that by using DeepMicroClass as a preliminary classification step on metagenomic/viromic assemblies, one can further focus on the interested sequence types for the following analysis, such as metagenomic binning of prokaryotic or eukaryotic contigs, comparative genomic analysis of viral or plasmid sequences, etc. We conclude DeepMicroClass achieves higher performance than the other benchmarked predictors, and its application can facilitate studies of under-appreciated sequence types, such as microbial eukaryotic or viral sequences.

## Availability of data and materials

The source code and user guide are available at https://github.com/chengsly/DeepMicrobeFinder. Benchmark datasets have been deposited at figshare (available at dx.doi.org/10.6084/m9.figshare.14576193). Raw reads for the case study were deposited at NCBI under the umbrella bioproject PRJNA739254.

Additional details of data and analysis are available from the corresponding authors upon request.

## Authors’ contributions

SH, JAF, and FS conceived the project; SH, TT, SC and FS designed the neural network structure and model evaluation procedures; SH and TT designed the training, test datasets and use-case applications; SH, TT and SC prepared the training and test datasets; TT, SC and SH implemented the software and performed the data analysis; SH, TT and SC prepared all the figures and tables; SH drafted the manuscript; TT, SC, TC, JAF and FS reviewed and edited the manuscript.

## Acknowledgements

This study was supported by the NIH grant (Grant ID: 2125142) to F. Sun, the Simons Collaboration on Computational Biogeochemical Modeling of Marine Ecosystems/CBIOMES) grant (Grant ID: 549943) and the Gordon and Betty Moore Foundation (Grant Number: 3779) to J. Fuhrman, the NSFC grants to S. Hou (Grant ID: 42276163) and T. Chen (Grant ID: 61872218, 61721003), the Shenzhen Science, Technology and Innovation Commission Programme to S. Hou (Grant ID: JCYJ20220530115401003), and the National Key R&D Program of China (Grant ID: 2019YFB1404804) to T. Chen. The funders had no roles in study design, data collection or analysis, the decision to publish, and the preparation of the manuscript. We thank Dr. David M. Needham, Dr. J. Cesar Ignacio-Espinoza, and Erin B. Fichot for their help with DNA extraction and metagenomic library preparation.

## List of abbreviations

Abbreviations used in this manuscript:

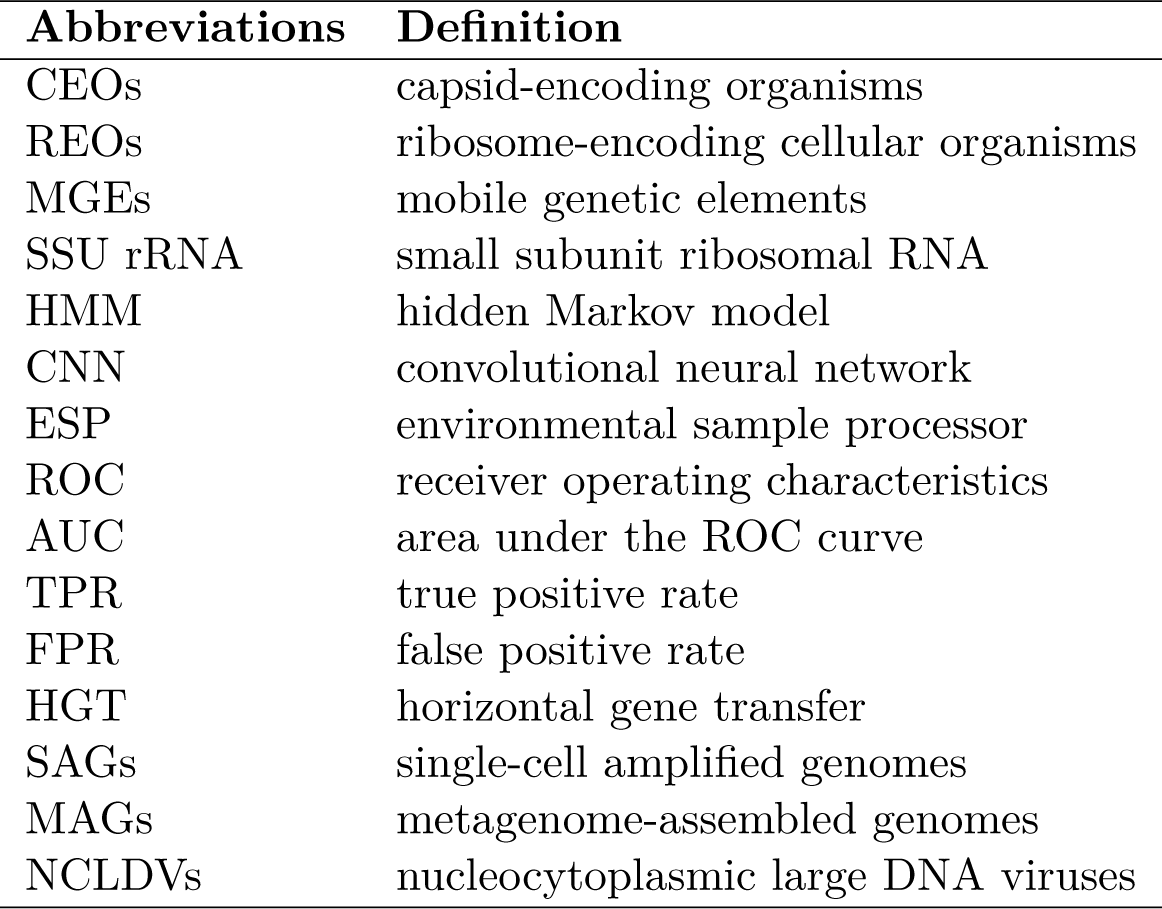

## Supporting information

Supplemental Table S1. The composition of 20 benchmark datasets used in this study.

PROK includes prokaryotic genomes, plasmids and prokaryotic viruses; EUK includes eukaryotic genomes and viruses. Prok: prokaryotic genomes, ProkVir: prokaryotic viruses/phages, Plas: plasmids, Euk: eukaryotic genomes, EukVir: eukaryotic viruses. Benchmark sequence files can be found at dx.doi.org/10.6084/m9.figshare.14576193.

Supplemental Figure S1. **Sequence source composition of 20 equal-sized benchmark datasets.** The fractions of PROK (including prokaryotic hosts, prokaryotic viruses, and plasmids) to EUK (including eukaryotic hosts and eukaryotic viruses) sequences were determined using the ratios of 9:1, 7:3, 5:5, 3:7, and 1:9. For each fixed PROK:EUK ratio, the PROK fraction was further split into prokaryotic hosts, prokaryotic viruses and plasmids based on the ratios of 5:1:1, 4:1:1, 3:1:1, and 2:1:1; and the EUK fraction was further split into eukaryotic hosts and eukaryotic viruses according to the ratio of 5:1, 4:1, 3:1, and 2:1. The detailed ratios can be found in Table S1.

Supplemental Figure S2. The distribution of viral confidence scores for (a) VirFinder and **(b) PPR-Meta. For both predictors, the same dataset was used and the predictions were performed with default parameters.** VirFinder uses VF-Scores to determine the likelihood of input sequences being viral or not, and PPR-Meta uses phage scores to discern viruses from host chromosomes and plasmids. Both predictors achieved a high recall for prokaryotic viruses, while the confidence scores of eukaryotic viruses were more evenly spread across all confidence regions. Besides, both predictors achieved a high performance in distinguishing prokaryotic host sequences from prokaryotic viruses, but less so for eukaryotic host sequences.

Supplemental Figure S3. **Performance of DeepMicroClass, Tiara and Whokaryote on eukaryotic sequence classification.** Both the accuracy and F1 score were compared based on 20 designed benchmark datasets. The sequence class composition of the 20 datasets can be found in Table S1. Values on top of the pairwise comparisons are Bonferroni adjusted t-test *p*-values.The significance of the overall ANOVA test was shown in the bottom left corner.

Supplemental Figure S4. **The distribution of misclassified sequence types by Tiara and Whokaryote.** The distribution of misclassified sequence types by Tiara and Whokaryote. The sequence composition of these datasets can be found in Table Supplemental Table S1. To make the figure more visible, the range of the *y*-axis is from 0 to 100 for Tiara and from 0 to 500 for Whokaryote.

Supplemental Figure S5. **Performance of DeepMicroClass, PlasFlow, PPR-Meta and geNomad on plasmid sequence classification.** Both the accuracy and F1 score were compared based on 20 designed benchmark datasets. The sequence class composition of the datasets can be found in Table S1. Values on top of the pairwise comparisons are Bonferroni-adjusted t-test *p*-values. The significance of the overall ANOVA test is shown in the bottom left corner.

Supplemental Figure S6. **The distribution of misclassified sequence types by PlasFlow, PPR-Meta and geNomad.** The sequence composition of these datasets can be found in Table S1.

Supplemental Figure S7. **Performance of DeepMicroClass (DMC), DeepVirFinder (DVF), VIBRANT, PPR-Meta (PPR) and geNomad on prokaryotic viral sequence classification.** Both the accuracy and F1 score were compared based on 20 designed benchmark datasets. The sequence class composition of the 20 test datasets can be found in Table S1. Values on top of the pairwise comparisons are Bonferroni-adjusted t-test *p*-values. The significance of the overall ANOVA test is shown in the bottom left corner

Supplemental Figure S8. **Performance of DeepMicroClass and VirSorter2 on prokaryotic and eukaryotic viral sequence classification.** Both the accuracy and F1 score were compared based on 20 designed benchmark datasets. The sequence class composition of these datasets can be found in Table S1. Values on top of the pairwise comparisons are Bonferroni-adjusted t-test *p*-values. The significance of the overall ANOVA test is shown in the bottom left corner.

Supplemental Figure S9. **TThe distribution of misclassified sequence types by PPR-Meta, DeepVirFinder, VIBRANT, geNomad and VirSorter2.** For PPR-Meta, DeepVirFinder, VIBRANT and geNomad, only prokaryotic viruses are considered as the positive set, and for VirSorter2 both prokaryotic and eukaryotic viruses are considered positive. The sequence composition of these datasets can be found in Table S1. To make the figure more visible, the range of the *y*-axis is from 0 to 500 for PPR-Meta and DeepVirFinder, from 0 to 50 for VIBRANT, and from 0 to 80 for VirSorter2.

Supplemental Figure S10. **Distribution patterns of accuracy (a) and F1 score (b) across 20 benchmark datasets for DeepMicroClass, PPR-Meta and geNomad on the prokaryotic genome, prokaryotic virus and plasmid classification.** DeepMicroClass received higher scores in both accuracy and F1 score metrics in all tested scenarios compared to PPR-Meta and geNomad in multi-class classification. The dashed black lines indicate where accuracy or F1 score equals 0.8. The same benchmark datasets were used as in Fig. **Fig. 3**.

Supplemental Figure S11. **Performance of DeepMicroClass, PPR-Meta and geNomad on the prokaryotic genome, prokaryotic virus and plasmid classification.** Both the accuracy and F1 score were compared based on 20 designed benchmark datasets. The sequence class composition of these datasets can be found in Table S1. Values on top of the pairwise comparisons are Bonferroni-adjusted t-test *p*-values. The significance of the overall ANOVA test is shown in the bottom left corner.

Supplemental Figure S12. **The distribution of misclassified sequence types by DeepMicroClass.** The sequence composition of these datasets can be found in Table S1. The maximal number of errors across all benchmark datasets was 50, which was set as the maximum of the *y*-axis.

Supplemental Figure S13. **Correlation coefficients of Prokaryotic (a), Eukaryotic (b), ProkaryoticViral (c), and EukaryoticViral (d) sequence relative abundances of different sequence classifiers.** Coefficients highlighted in colors are significant ones (*p*-value *<* 0.01).

