## Supplemental Information for "DeepMicroClass sorts metagenomes into prokaryotes, eukaryotes and viruses, with marine applications"

### Supplementary Materials

#### Supplemental Tables

| Dataset_No | Prok | ProkVir | Plas | Euk | EukVir | PROK | EUK | PROK :EUK | Prok:ProkVir:Plas \| Euk:EukVir |
|---|---|---|---|---|---|---|---|---|---|
| DS_1 | 643 | 129 | 129 | 83 | 17 | 901 | 100 | 9:1 | 5:1:1 \| 5:1 |
| DS_2 | 600 | 150 | 150 | 80 | 20 | 900 | 100 | 9:1 | 4:1:1 \| 4:1 |
| DS_3 | 540 | 180 | 180 | 75 | 25 | 900 | 100 | 9:1 | 3:1:1 \| 3:1 |
| DS_4 | 450 | 225 | 225 | 67 | 33 | 900 | 100 | 9:1 | 2:1:1 \| 2:1 |
| DS_5 | 500 | 100 | 100 | 250 | 50 | 700 | 300 | 7:3 | 5:1:1 \| 5:1 |
| DS_6 | 467 | 117 | 117 | 240 | 60 | 701 | 300 | 7:3 | 4:1:1 \| 4:1 |
| DS_7 | 420 | 140 | 140 | 225 | 75 | 700 | 300 | 7:3 | 3:1:1 \| 3:1 |
| DS_8 | 350 | 175 | 175 | 200 | 100 | 700 | 300 | 7:3 | 2:1:1 \| 2:1 |
| DS_9 | 357 | 71 | 71 | 417 | 83 | 499 | 500 | 5:5 | 5:1:1 \| 5:1 |
| DS_10 | 333 | 83 | 83 | 400 | 100 | 499 | 500 | 5:5 | 4:1:1 \| 4:1 |
| DS_11 | 300 | 100 | 100 | 375 | 125 | 500 | 500 | 5:5 | 3:1:1 \| 3:1 |
| DS_12 | 250 | 125 | 125 | 333 | 167 | 500 | 500 | 5:5 | 2:1:1 \| 2:1 |
| DS_13 | 214 | 43 | 43 | 583 | 117 | 300 | 700 | 3:7 | 5:1:1 \| 5:1 |
| DS_14 | 200 | 50 | 50 | 560 | 140 | 300 | 700 | 3:7 | 4:1:1 \| 4:1 |
| DS_15 | 180 | 60 | 60 | 525 | 175 | 300 | 700 | 3:7 | 3:1:1 \| 3:1 |
| DS_16 | 150 | 75 | 75 | 467 | 233 | 300 | 700 | 3:7 | 2:1:1 \| 2:1 |
| DS_17 | 71 | 14 | 14 | 750 | 150 | 99 | 900 | 1:9 | 5:1:1 \| 5:1 |
| DS_18 | 67 | 17 | 17 | 720 | 180 | 101 | 900 | 1:9 | 4:1:1 \| 4:1 |
| DS_19 | 60 | 20 | 20 | 675 | 225 | 100 | 900 | 1:9 | 3:1:1 \| 3:1 |
| DS_20 | 50 | 25 | 25 | 600 | 300 | 100 | 900 | 1:9 | 2:1:1 \| 2:1 |

**Table S1.** The composition of 20 benchmark datasets used in this study. PROK includes prokaryotic genomes, plasmids and prokaryotic viruses; EUK includes eukaryotic genomes and viruses. Prok: prokaryotic genomes, ProkVir: prokaryotic viruses/phages, Plas: plasmids, Euk: eukaryotic genomes, EukVir: eukaryotic viruses. Benchmark sequence files can be found at dx.doi.org/10.6084/m9.figshare.14576193.

### Supplemental Figures

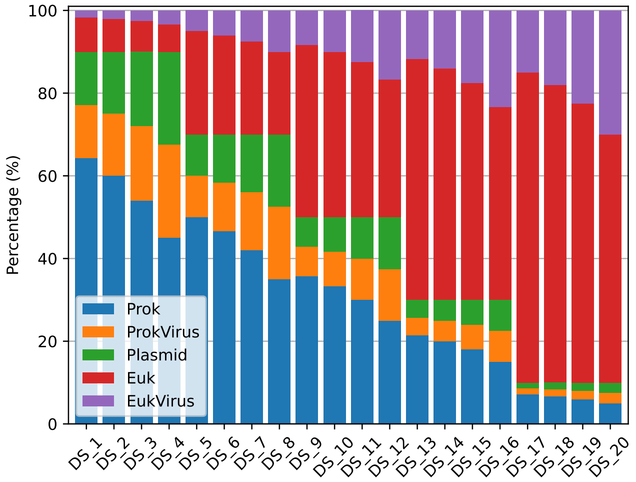


**Fig S1. Sequence source composition of 20 equal-sized benchmark datasets.** The fractions of PROK (including prokaryotic hosts, prokaryotic viruses, and plasmids) to EUK (including eukaryotic hosts and eukaryotic viruses) sequences were determined using the ratios of 9:1, 7:3, 5:5, 3:7, and 1:9. For each fixed PROK:EUK ratio, the PROK fraction was further split into prokaryotic hosts, prokaryotic viruses and plasmids based on the ratios of 5:1:1, 4:1:1, 3:1:1, and 2:1:1; and the EUK fraction was further split into eukaryotic hosts and eukaryotic viruses according to the ratio of 5:1, 4:1, 3:1, and 2:1. The detailed ratios can be found in Table S1.

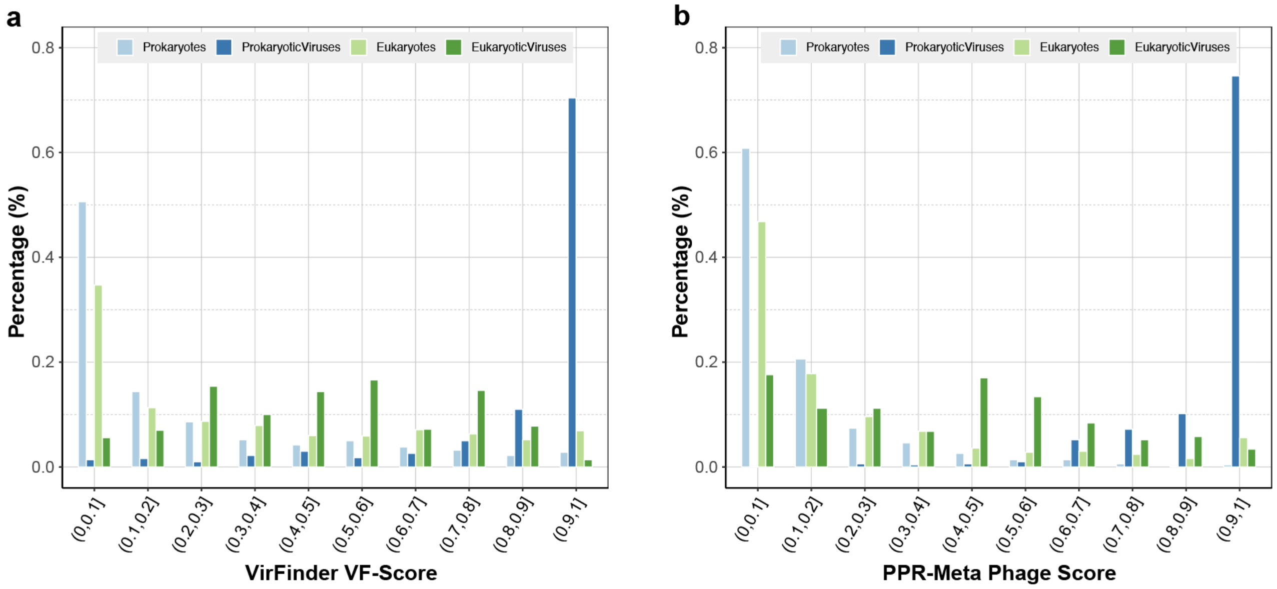


**Fig S2. The distribution of viral confidence scores for (a) VirFinder and (b) PPR-Meta.** For both predictors, the same dataset was used and the predictions were performed with default parameters. VirFinder uses VF-Scores to determine the likelihood of input sequences being viral or not, and PPR-Meta uses phage scores to discern viruses from host chromosomes and plasmids. Both predictors achieved a high recall for prokaryotic viruses, while the confidence scores of eukaryotic viruses were more evenly spread across all confidence regions. Besides, both predictors achieved a high performance in distinguishing prokaryotic host sequences from prokaryotic viruses, but less so for eukaryotic host sequences.

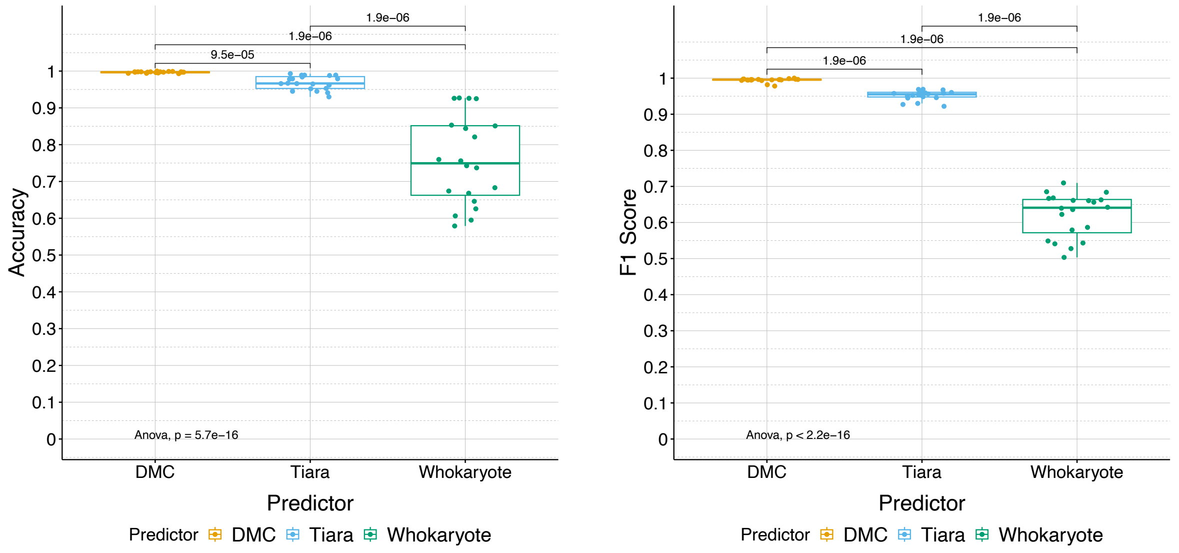


**Fig S3. Performance of DeepMicroClass, Tiara and Whokaryote on eukaryotic sequence classification.** Both the accuracy and F1 score were compared based on 20 designed benchmark datasets. The sequence class composition of the 20 datasets can be found in Table S1. Values on top of the pairwise comparisons are Bonferroni adjusted t-test $p$-values.The significance of the overall ANOVA test was shown in the bottom left corner.

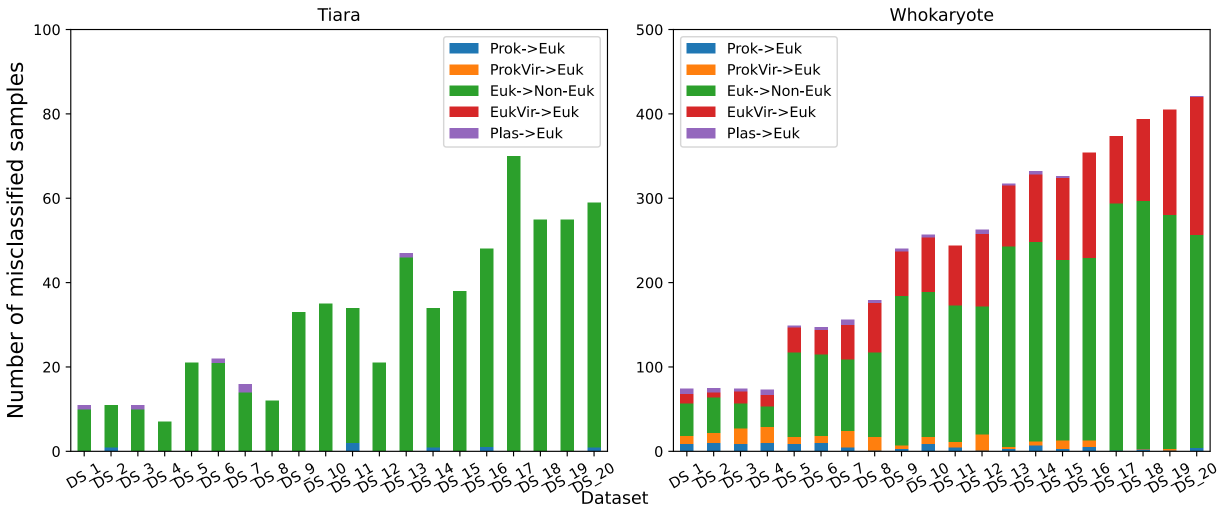


**Fig S4. The distribution of misclassified sequence types by Tiara and Whokaryote.** The sequence composition of these datasets can be found in Table S1. To make the figure more visible, the range of the $y$-axis is from 0 to 100 for Tiara and from 0 to 500 for Whokaryote.

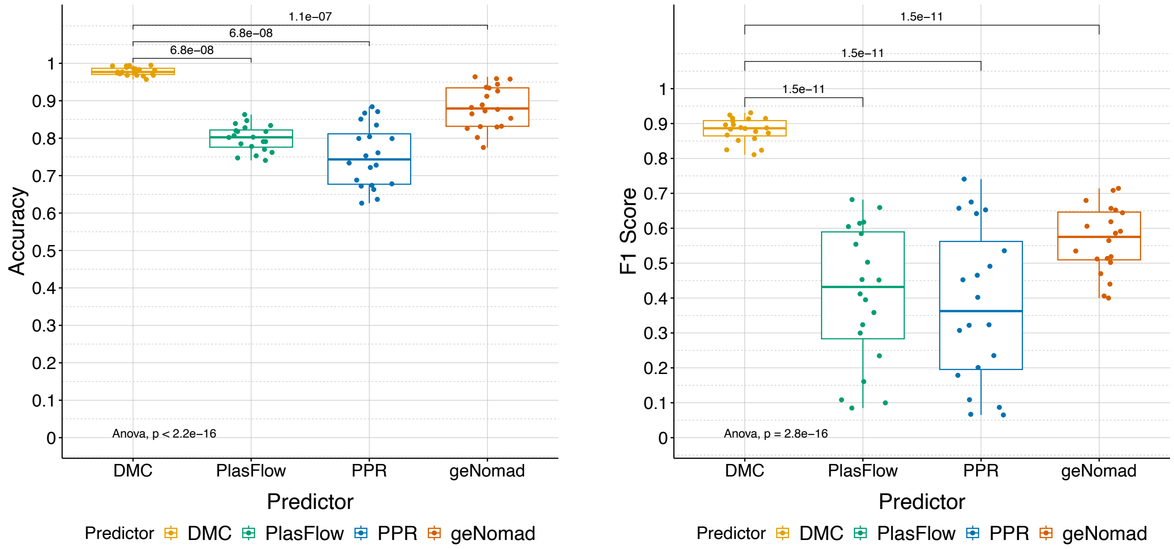


**Fig S5. Performance of DeepMicroClass, PlasFlow, PPR-Meta and geNomad on plasmid sequence classification.** Both the accuracy and F1 score were compared based on 20 designed benchmark datasets. The sequence class composition of the datasets can be found in Table S1. Values on top of the pairwise comparisons are Bonferroni adjusted t-test $p$-values. The significance of the overall ANOVA test is shown in the bottom left corner.

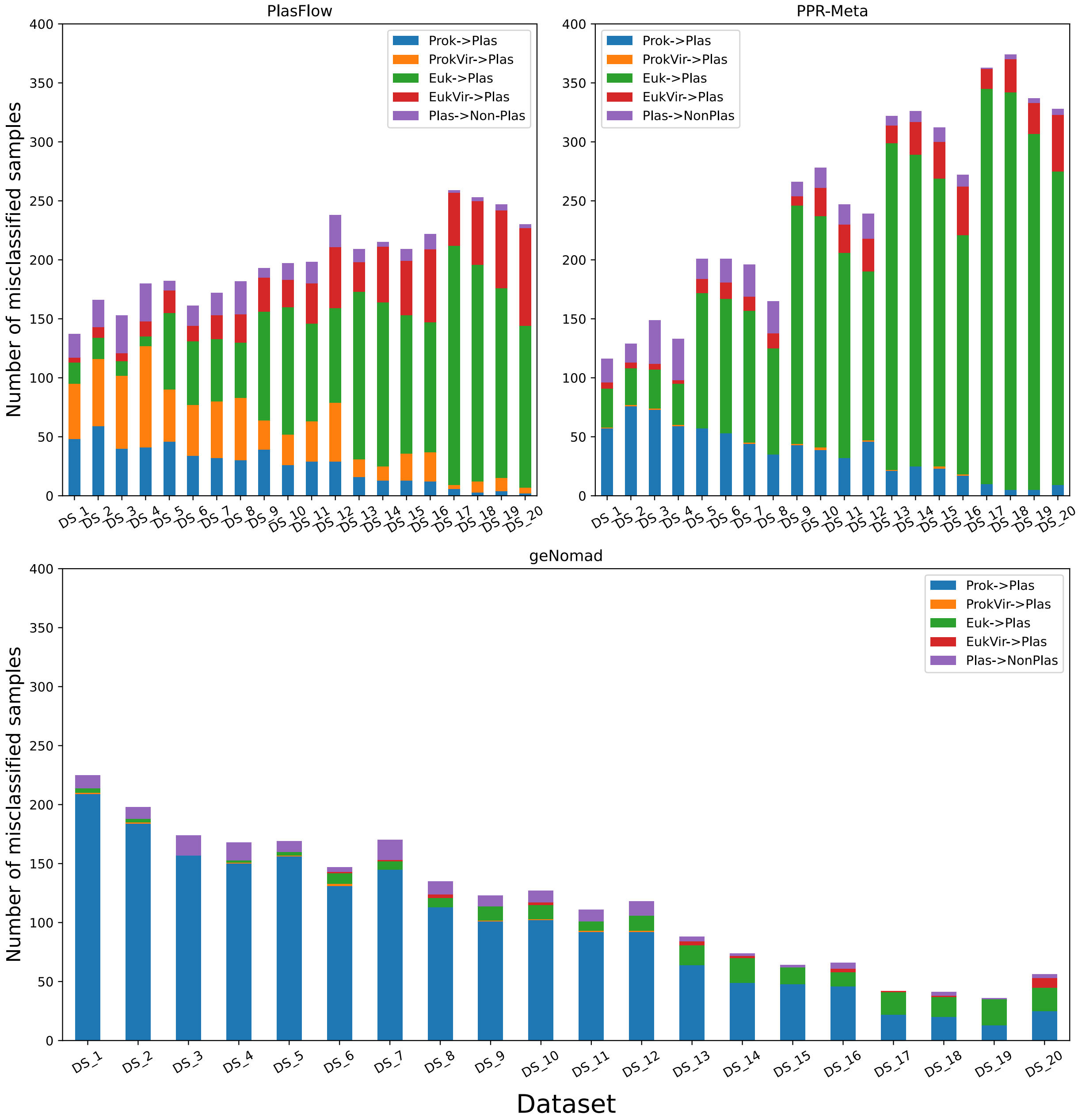


**Fig S6. The distribution of misclassified sequence types by PlasFlow, PPR-Meta and geNomad.** The sequence composition of these datasets can be found in Table S1. The y-axis ranges from 0 to 400 for both panels.

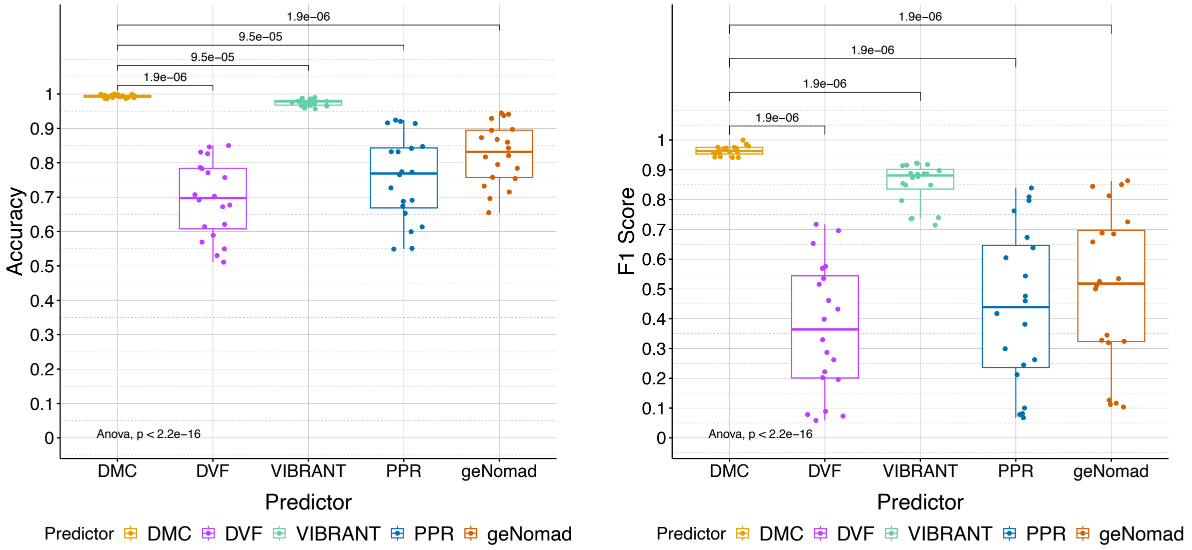


**Fig S7. Performance of DeepMicroClass (DMC), DeepVirFinder (DVF), VIBRANT, PPR-Meta (PPR) and geNomad on prokaryotic viral sequence classification.** Both the accuracy and F1 score were compared based on 20 designed benchmark datasets. The sequence class composition of these datasets can be found in Table S1. Values on top of the pairwise comparisons are Bonferroni adjusted t-test $p$-values. The significance of the overall ANOVA test is shown in the bottom left corner.

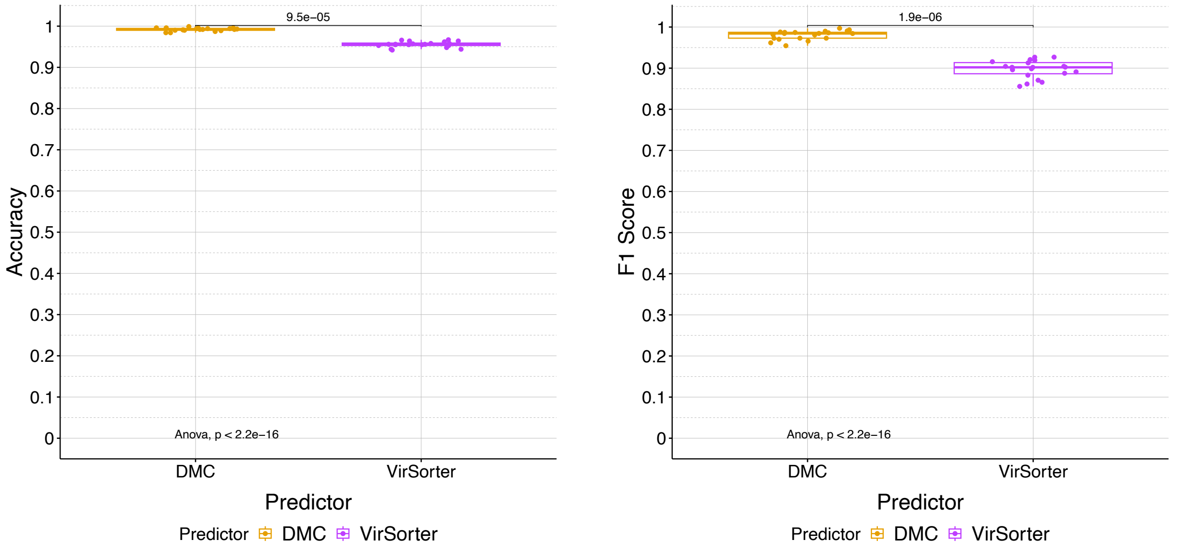


**Fig S8. Performance of DeepMicroClass and VirSorter2 on prokaryotic and eukaryotic viral sequence classification.** Both the accuracy and F1 score were compared based on 20 designed benchmark datasets. The sequence class composition of these datasets can be found in Table S1. Values on top of the pairwise comparisons are Bonferroni adjusted t-test $p$-values. The significance of the overall ANOVA test is shown in the bottom left corner.

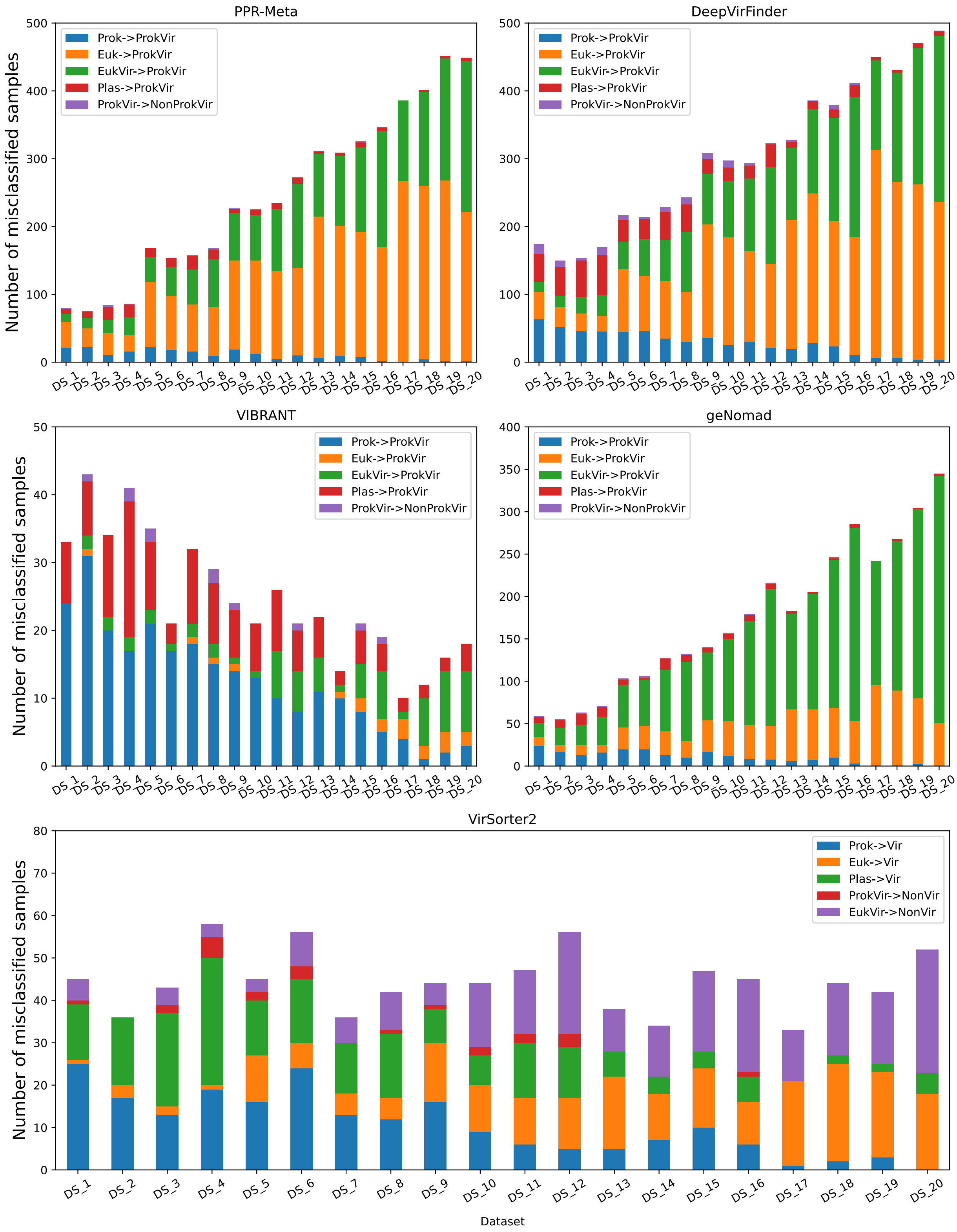


**Fig S9. The distribution of misclassified sequence types by PPR-Meta, DeepVirFinder, VIBRANT, geNomad and VirSorter2.** For PPR-Meta, DeepVirFinder, VIBRANT and geNomad, only prokaryotic viruses are considered as the positive set, and for VirSorter2 both prokaryotic and eukaryotic viruses are considered positive. The sequence composition of these datasets can be found in Table S1. To make the figure more visible, the range of y-axis is from 0 to 500 for PPR-Meta and DeepVirFinder, from 0 to 50 for VIBRANT, and from 0 to 80 for VirSorter2.

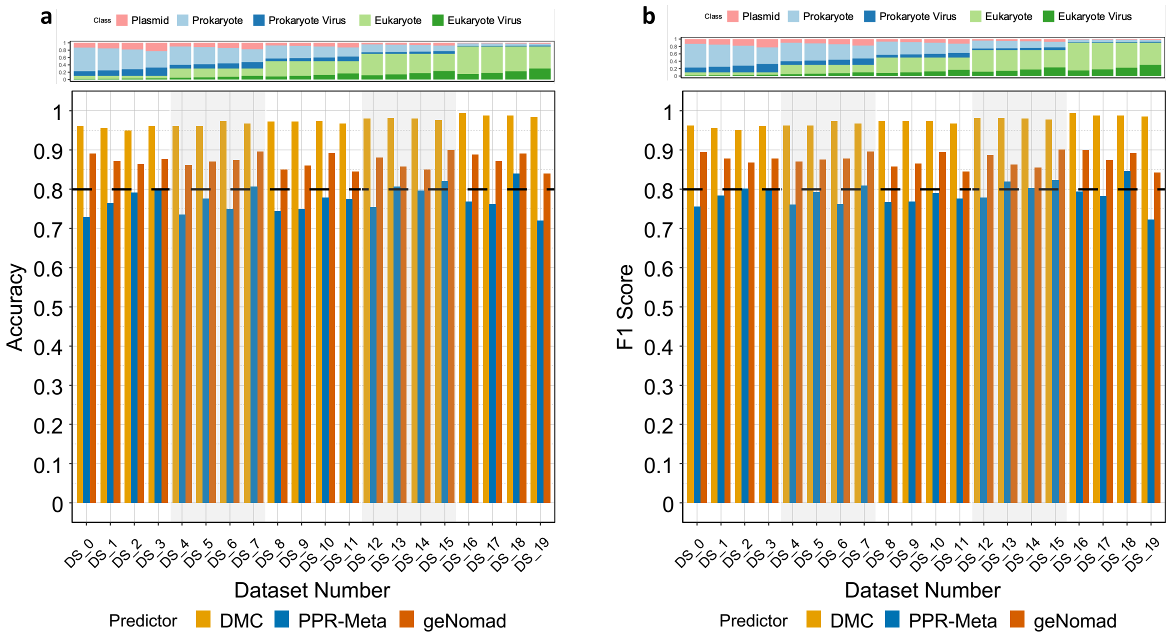


**Fig S10. Distribution patterns of accuracy (a) and F1 score (b) across 20 benchmark datasets for DeepMicroClass, PPR-Meta and geNomad on the prokaryotic genome, prokaryotic virus and plasmid classification.** DeepMicroClass received higher scores in both accuracy and F1 score metrics in all tested scenarios compared to PPR-Meta and geNomad in multi-class classification. The dashed black lines indicate where accuracy or F1 score equals 0.8. The same benchmark datasets were used as in **Fig. 3**.

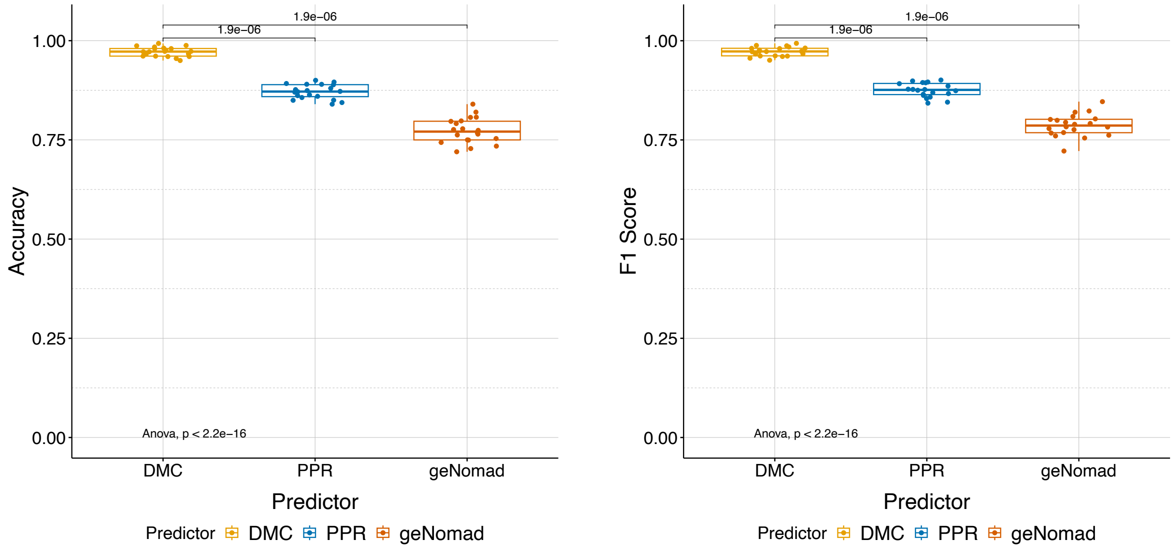


**Fig S11. Performance of DeepMicroClass, PPR-Meta and geNomad on the prokaryotic genome, prokaryotic virus and plasmid classification.** Both the accuracy and F1 score were compared based on 20 designed benchmark datasets. The sequence class composition of these datasets can be found in Table S1. Values on top of the pairwise comparisons are Bonferroni adjusted t-test $p$-values. The significance of the overall ANOVA test is shown in the bottom left corner.

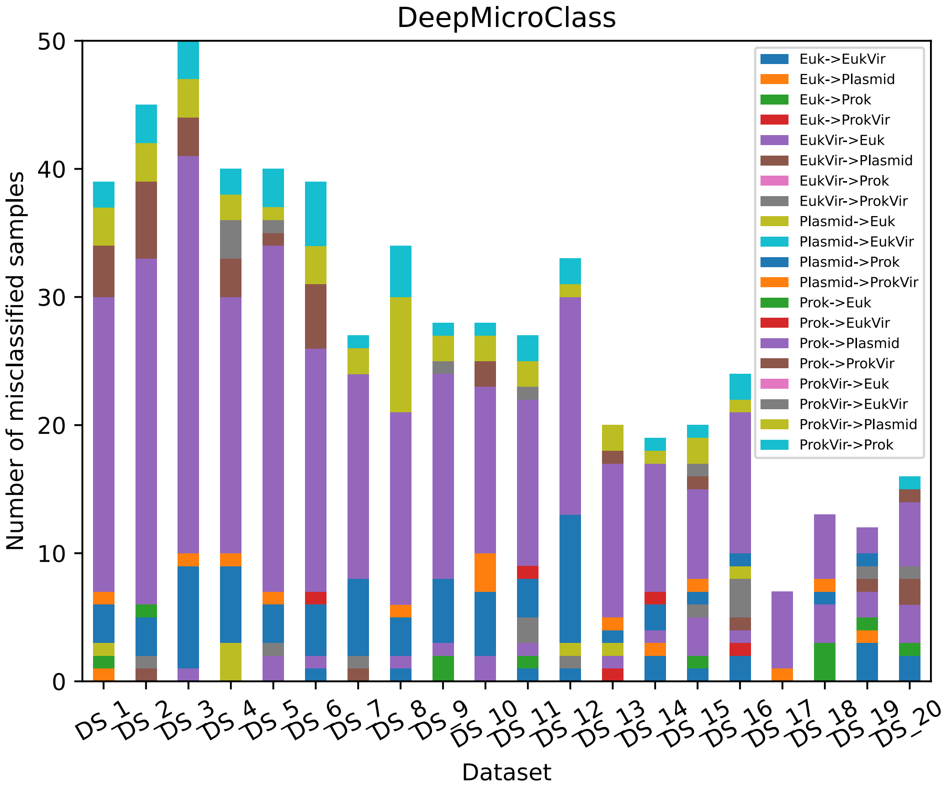


**Fig S12. The distribution of misclassified sequence types by DeepMicroClass.** The sequence composition of these datasets can be found in Table S1. The maximal number of errors across all benchmark datasets was 50, which was set as the maximum of the y-axis.

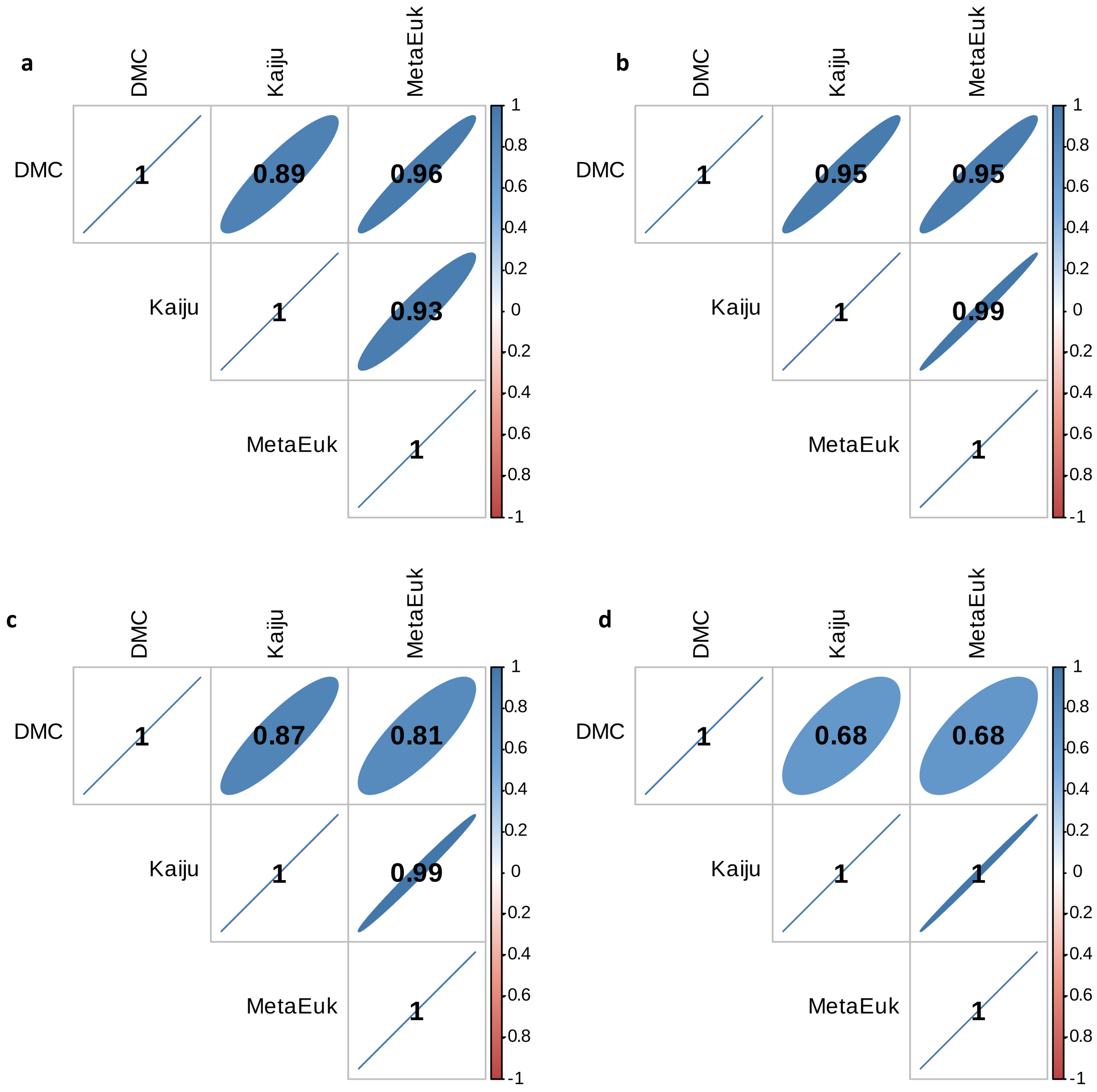


**Fig S13. Correlation coefficients of Prokaryotic (a), Eukaryotic (b), ProkaryoticViral (c), and EukaryoticViral (d) sequence relative abundances of different sequence classifiers.** Coefficients highlighted in colors are significant ones ($p$-value $< 0.01$).
